# Voltage tunes mGlu5 receptor function, impacting synaptic transmission

**DOI:** 10.1101/2023.07.17.549279

**Authors:** Marin Boutonnet, Camille Carpena, Nathalie Bouquier, Yan Chastagnier, Joan Font-Ingles, Enora Moutin, Ludovic Tricoire, Jean Chemin, Julie Perroy

## Abstract

Voltage sensitivity is a common feature of many membrane proteins, including some G-protein coupled receptors (GPCRs). However, the functional consequences of voltage sensitivity in GPCRs are not well understood.

In this study, we investigated the voltage sensitivity of the post-synaptic metabotropic glutamate receptor mGlu5 and its impact on synaptic transmission. Using biosensors and electrophysiological recordings in non-excitable HEK293T cells or neurons, we found that mGlu5 receptor function is optimal at resting membrane potentials. We observed that membrane depolarization significantly reduced mGlu5 receptor activation, Gq-PLC/PKC stimulation, Ca^2+^ release, and mGlu5 receptor-gated currents through TRPC6 channels or NMDA receptors. Notably, we report a previously unknown activity of the NMDA receptor at the resting potential of neurons, enabled by mGlu5.

Our findings suggest that mGlu5 receptor activity is directly regulated by membrane voltage which may have a significant impact on synaptic processes and pathophysiological functions.

## Introduction

G-protein coupled receptors (GPCRs) represent a class of 7-transmembrane receptors that are involved in numerous physiological processes and remain the most extensively targeted protein family by approved drugs (1). Recent research has unveiled that the activity of certain GPCRs can be modulated by the membrane voltage (V_m_). In neurons, where V_m_ undergoes permanent changes, this emerging property could significantly impact the functioning of neurotransmitter-activated GPCRs and their role as modulators of synaptic transmission. Indeed, V_m_ has been shown to positively or negatively regulate the function of several GPCRs, about thirty out of the thousand members of this big family, including acetylcholine (2–5), purine (6), opioid (7), dopamine (8), and prostanoid receptors (9). Although the structural mechanisms that underlie the sensitivity of GPCRs to V_m_ are still being elucidated (4,10–13), functional studies using site-directed mutagenesis of the V_m_ sensor suggest that GPCR activity can be affected, leading to impaired neurotransmitter release (14,15) or synaptic plasticity and behavior (16).

Glutamate is the primary excitatory neurotransmitter in the brain that binds to both ionotropic (AMPA, NMDA, Kainate receptors) and metabotropic glutamate (mGlu) receptors. The mGlu receptor family, comprising eight G protein-coupled receptors (mGlu1-8), has been extensively studied for their modulatory role in synaptic transmission and plasticity (17). Preclinical and clinical studies have targeted these receptors in various neurological disorders, such as Autism, Fragile X syndrom, Schizophrenia, Parkinson’s, and Alzheimer’s disease (18). So far, only one study, in the Xenopus oocytes expression system, suggests a direct sensitivity of some mGlu receptors to V_m_ (mGlu1 and 3, (11)). Of note, membrane potential changes seem to play a synergistic role in neurons on signaling events mediated by mGlu5 (19) and mGlu7 (20) receptors, but without any evidence of a direct action of V_m_ on the receptor itself. Yet, due to their synaptic location, these receptors are permanently exposed to membrane voltage fluctuations. Therefore, demonstrating the sensitivity of mGlu receptors to V_m_ could provide insights into their role depending on the state of neuronal activity. In this study, we have selected the postsynaptic mGlu5 receptor as a model receptor, known for its role as neuromodulator of synaptic transmission. mGlu5 is also a key trigger in the induction of synaptic plasticity, working in concert with ionotropic NMDA receptors through physical and functional crosstalk (21). Interestingly, NMDA receptors are a prototype of receptors whose activity is regulated by the membrane potential. As a detector of neuronal activity coincidence, NMDA activation is limited to synapses whose pre- and postsynaptic elements are activated simultaneously (22,23). Thus, a sensitivity of the mGlu5 receptor to V_m_ could similarly specify the identity of synapses on which mGlu5 exerts its functional effects depending on the neuron’s activity history.

In this report, we explore the intricate signaling pathways associated with the conformational change of postsynaptic mGlu5 receptor induced by glutamate binding. This conformational change leads to the activation of G_q/11_-type proteins, the activation of the PLC-PKC pathway, and the subsequent release of Ca^2+^ from internal stores through IP3 receptors (18), ultimately fine-tuning synaptic transmission by gating channels and ionotropic receptors, including NMDA (24–26), AMPA (27,28), and transient receptor potential canonical (TRPC) channels (29). Specifically, we investigate the impact of V_m_ on each of the canonical signaling events of the mGlu5 receptor using a variety of biosensors and patch clamp recordings. Our results demonstrate that V_m_ modulates mGlu5 receptor activation and its downstream signaling, with a depolarization of the membrane favoring the inactive conformation of mGlu5 and leading to a decrease in mGlu5-induced G_q/11_ activation, Ca^2+^ signaling, TRPC6 gating, and NMDA receptor facilitation. Interestingly, our data reveals that the mGlu5 receptor functions optimally at potentials close to the resting potential of the cells.

## Results

### V_m_ regulates mGlu5-induced Ca^2+^ release

To investigate the effect of V_m_ on mGlu5 function and determine the optimal experimental conditions, we examined mGlu5-induced Ca^2+^ release from intracellular stores. mGlu5 stimulation is known to induce a transient increase of Ca^2+^ oscillations in various cell types, including cultured astrocytes (30), hippocampal neurons (19), midbrain neurons (31), as well as in heterologous expression systems such as Chinese Hamster Ovary cells (32) and Human Embryonic Kidney 293T (HEK293T) cells (33). In line with these findings, in HEK293T cells expressing the GCaMP6s fluorescent Ca^2+^ sensor (**Figure 1a**), mGlu5 agonist (RS)-3,5-dihydroxyphenylglycine (DHPG, 100 µM) induced a Ca^2+^ increase exclusively in cells expressing mGlu5, as evidenced by GCaMP6s fluorescence fluctuations measured in the cell population using a fluorimeter (**Figure 1b**). The mGlu5 agonist-induced Ca^2+^ increase in mGlu5-expressing cells was inhibited by application of MPEP (10 µM), a mGlu5-specific negative allosteric modulator (NAM), confirming the requirement for mGlu5 receptor activation (**Figure 1b**).

**Figure 1.**
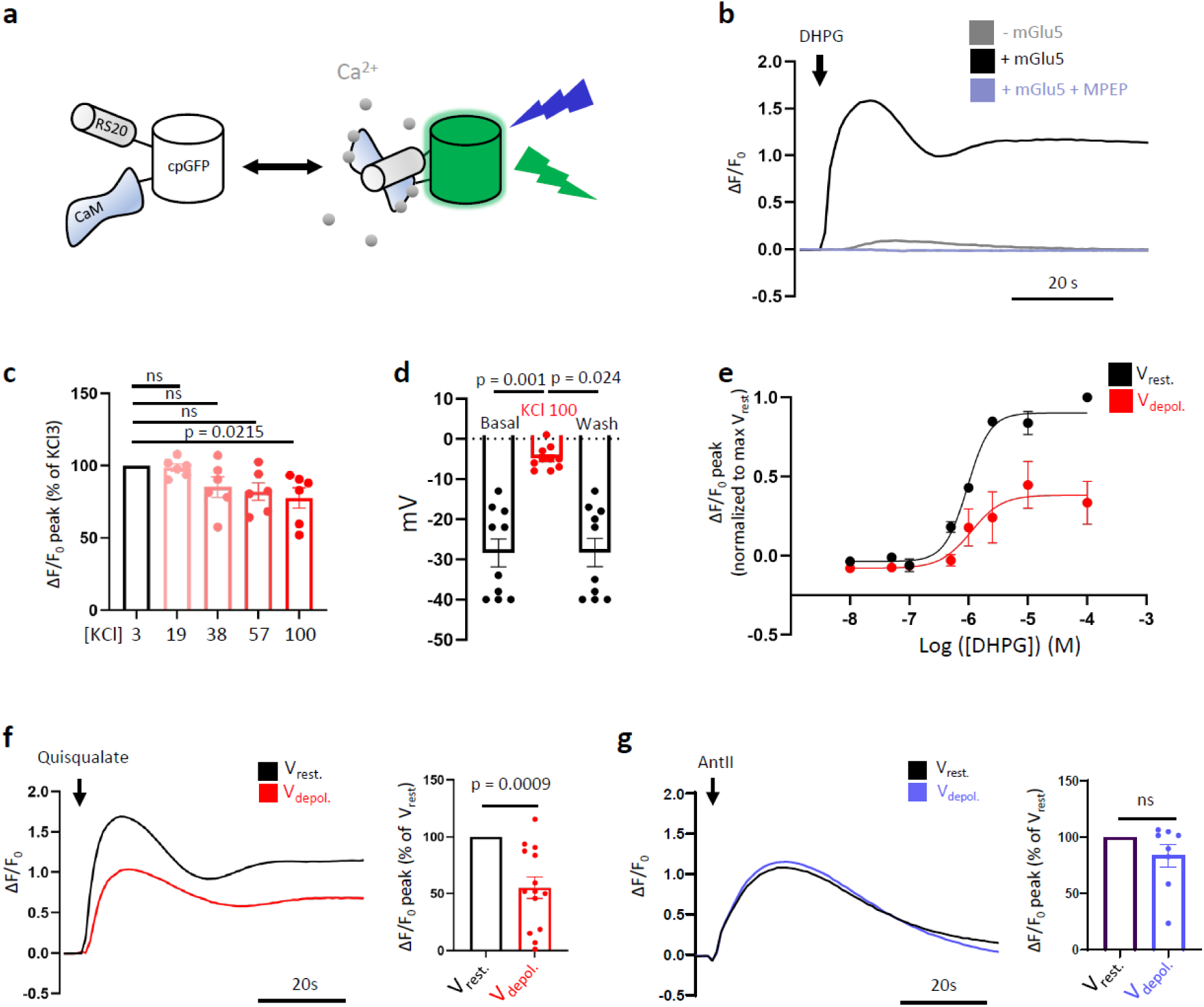
Voltage regulates mGlu5 agonist-induced Ca^2+^ release. HEK cells stably expressing GCaMP6s also transiently expressed mGlu5 (b-f) or AT1 (g) receptors. **a)** Scheme of GCaMP6s fluorescent Ca^2+^ sensor - adapted from (80). GCaMP6s is composed of a circularly permuted GFP, a calmodulin (CaM) and a peptide from smooth-muscle myosin light-chain kinase (RS20). **b)** DHPG (100µM)-induced fluorescence fluctuation in cells expressing (black and blue) or not (grey) mGlu5 receptor, in absence (grey and black) or presence (blue) of the mGlu5 NAM, MPEP (10µM); Representative illustration. **c)** Peak of fluorescence fluctuation induced by quisqualate (10µM) in isotonic extracellular solutions containing increasing KCl concentrations ([KCl] in mM), normalized to fluorescence fluctuations recorded with the 3mM [KCl] solution. Data are mean ± SEM of n = 6 triplicates from independent experiments. Statistics: Kruskal-Wallis test, **d)** Membrane potential measured in the current clamp whole-cell configuration with extracellular solutions containing either 3mM (black) or 100mM (red) KCl. Data are mean ± SEM of n = 10 cells from 3 independent experiments. Statistics: Friedman test **e)** Dose-response curves of DHPG-induced fluorescence fluctuation with KCl 3mM (black, V_rest_) and 100mM (red, V_depol_), normalized to V_rest_ peak of each experiment. Data are mean ± SEM of n = 3 independent experiments. **f** and **g)** Quisqualate (10µM, f) or Angiotensine II (1µM, g) - induced fluorescence fluctuation in KCl 3mM (V_rest_) and 100mM (V_depol_) solutions. Left, representative illustration; Right, mean ± SEM of n = 7 to 14 triplicates from independent experiments; Statistics: one sample Wilcoxon test.

In order to determine the effect of V_m_ on mGlu5-induced Ca^2+^ increase, we treated cells with equimolar solutions containing increasing concentrations of KCl (from 3 mM, a physiological concentration referred to as “V_rest_”, up to 100 mM) to induce a systematic, well-controlled, and long-lasting membrane depolarization (as previously demonstrated in neurons (34)). Interestingly, KCl 100 mM had no effect on basal intracellular Ca^2+^ concentrations (**Figure Supp. 1a**), but significantly reduced mGlu5 agonist-induced Ca^2+^ increase by 22.3 ± 7.1% compared to KCl 3 mM (**Figure 1c**). We measured that KCl 100 mM application increased the membrane potential by 23.6 ± 0.9 mV (**Figure 1d**), referred to as “V_depol_”. Whole cell patch clamp recordings indeed reported a reversible switch from V_rest_ = -28.4 ± 3.5 mV at physiological KCl concentration (3 mM) to V_depol_ = -4.8 ± 0.9 mV with KCl 100 mM (**Figure 1d**). The dose response curve of DHPG-induced Ca^2+^ release further revealed a reduction of the agonist efficacy at V_depol_ compared to V_rest_ (**Figure 1e**). Additionally, saturating concentrations of another mGlu5 agonist, quisqualate (10µM), triggered a Ca^2+^ increase at V_rest_, which was reduced by 45 ± 9.5% at V_depol_ (**Figure 1f**). These initial findings indicate that the ability of the mGlu5 receptor to increase intracellular Ca^2+^ is dependent on V_m_. To investigate whether the effect of V_m_ was specific to the mGlu5 receptor, we examined another G_q_-coupled receptor, the angiotensin II type 1 receptor (AT1), and did not observe any influence of V_m_ on AT1 receptor-mediated Ca^2+^ release, thereby supporting a specific effect of V_depol_ on mGlu5 signaling in this context (**Figure 1g**).

To investigate the effect of V_m_ on mGlu5-induced Ca^2+^ increase at a cellular level, we performed live cell imaging of GCaMP6s fluorescence (**Figure 2a** and **supplemental Video1**). Initially, large recordings in the entire field of view showed a global increase in Ca^2+^ induced by the mGlu5 agonist, quisqualate (10µM), at V_rest_. This increase was significantly reduced at V_depol_ (30 ± 10 % reduction compared to V_rest_ at the peak of fluorescence, **Figure 2b** and **supplemental Video2**). The global Ca^2+^ increase was an average of the single cell Ca^2+^ fluctuations, which were classified into 3 distinct populations of cells depending on the type of responses: silent cells, cells displaying 1 single Ca^2+^ spike, and Ca^2+^ oscillating cells (**Figure 2c**), as reported previously (35,36). V_depol_ modified the distribution of cells in each of these categories, with a global shift towards less oscillating populations (**Figure 2c**), and a significant reduction in the proportion of cells with an oscillating Ca^2+^ pattern (10.6% reduction of the proportion of oscillating cells compared to V_rest_, **Figure 2d**). Subsequently, for each oscillating cell, we studied the global frequency of Ca^2+^ oscillations induced by mGlu5 agonist. Quisqualate induced a mean of 0.49 ± 0.02 oscillations/min per cell at V_rest_, but only 0.39 ± 0.02 oscillations per cell at V_depol_. Thus, V_depol_ induced a 20.3 ± 4.8% reduction in the frequency of Ca^2+^ oscillations triggered by mGlu5 stimulation (**Figure 2e**).

**Figure 2.**
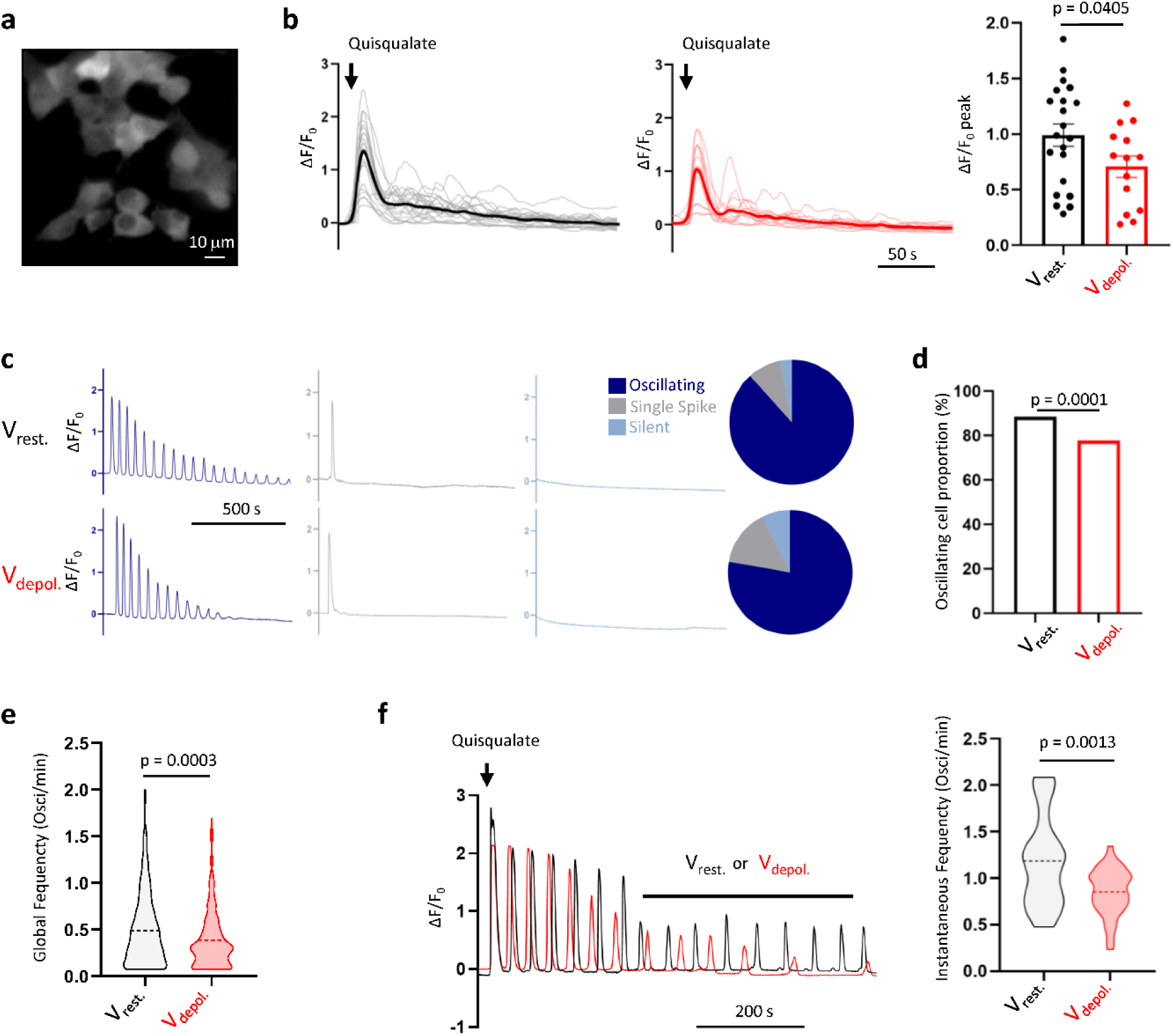
Voltage regulates mGlu5 agonist-induced Ca^2+^ oscillations. **a)** Representative field of GCaMP6s expressing cells in epifluorescence microscopy - Scale = 10 µm. **b)** Quisqualate (10µM)-induced fluorescence fluctuation at V_rest_ (left, black) and V_depol_ (middle, red), for individual field of view (thin) and average (thick) aligned to the peak of response, and quantification of the mean ± SEM of n = 14 to 21 independent experiments (right, Statistics: Mann-Whitney test) **c)** Fluorescence fluctuation recorded in representative oscillating, single spike and silent cells at V_rest_ and V_depol_ in response to quisqualate (10µM) with proportion of cells in each category displayed in a circular diagram, n = 403 cells for V_rest_ and 324 cells for V_depol_ **d)** Percentage of oscillating cells (> 2 oscillations) at V_rest_ and V_depol_. n = 324 to 403 cells from 14 to 18 independent experiments. Statistics: N-1 χ² test, p = 0.0001. **e)** Global frequency of fluorescence oscillations induced by quisqualate (10µM) application at V_rest_ and V_depol_. Data are mean ± SEM of n = 280 to 408 cells. Statistics: unpaired t-test **f)** Representative illustrations (left) and violin plots with mean indicated by a dotted line (right) of fluorescence oscillations frequency induced by V_rest_ or V_depol_ imposed after quisqualate (10µM) application (n = 21 to 35 cells - Statistics: unpaired t-test)

Given that depolarization in neurons can also occur following receptor stimulation, we investigated the impact of V_depol_ subsequent to mGlu5 activation. Specifically, we measured the effect of V_depol_ compared to V_rest_ on oscillating cells that had undergone Ca^2+^ oscillations at resting membrane potential as a result of mGlu5 stimulation. To do so, we determined the instantaneous frequency, which represents the average inter-oscillation frequency between the first and last oscillations. Our findings consistently revealed a significant reduction in the instantaneous frequency, from 1.18 ± 0.11 at V_rest_ to 0.85 ± 0.04 oscillation/min at V_depol_, representing a 28 ± 4.7% decrease (**Figure 2f**). These results suggest that a membrane depolarization of approximately 20-25mV, whether it occurs before or after mGlu5 activation, can diminish the receptor’s ability to induce Ca^2+^ release from intracellular stores, in few seconds. Specifically, V_depol_ reduced the number of agonist-responsive cells and the efficiency of the responding cells, resembling the characteristics of a NAM (35,36).

### V_m_ tunes mGlu5 receptor probability of activation

The reduced ability of mGlu5 receptor to induce Ca^2+^ release from intracellular stores at V_depol_ may be attributed to a decrease in the expression of receptors at the plasma membrane and/or a diminished activation of these receptors under depolarized conditions. To investigate the first hypothesis, we measured the receptor cell surface expression using an enhance bystander Bioluminescence Resonance Energy Transfer (ebBRET) biosensor. ebBRET is based on naturally interacting chromophores luciferase (RLuc) and green fluorescent protein (rGFP) from *Renilla reniformis*, enabling to quantify the cell surface expression of an Rluc-tagged protein expressed with the rGFP anchored at the plasma membrane by a CAAX sequence ((37) **Figure Supp. 2a**). In mGlu5-RlucII and rGFP-CAAX co-transfected cells, incubation of MPEP for 5 to 30 minutes resulted in an expected increase in receptor expression at the cell surface, as reported by an increase in ebBRET (**Figure Supp. 2b**). This control experiment validated the use of the ebBRET biosensor. However, we did not observe any ebBRET variation induced by shorter (2 min) applications of MPEP or quisqualate, nor by V_depol_ alone or combined with these ligands (**Figure Supp. 2c**). Therefore, this experiment ruled out any potential effect of V_depol_ on the receptor cell surface expression, in these experimental conditions.

To investigate the proportion of active and inactive mGlu5 receptors at the cell surface, we then used a Time-Resolved Fluorescence Resonance Energy Transfer (TR-FRET) biosensor, reporting structural dynamics and drug action at the mGlu5 receptor (38,39). The conformational changes of the extracellular ligand-binding domains (ECDs) of mGlu5 dimers are associated with receptor activation: SNAP-mGlu5 (ST-mGlu5) labeled with the Lumi4-Tb donor and the fluorescein acceptor, display a high FRET signal in the absence of ligand or in presence of the antagonist and a low FRET signal in the presence of glutamate or other orthosteric agonists (**Figure 3a**). Any approach expected to stabilize the active conformation of the effector domain increases the agonist potency in stabilizing the active ECDs conformation (38). Hence, TR-FRET measurements allowed to determine the proportion of active and inactive ST-mGlu5 receptors, depending on V_m_. Importantly, labelling mGlu5 receptors separately on distinct cell populations, we first controlled that neither the fluorescein nor the Lumi4-Tb emissions were affected by V_depol_ (**Figure Supp 1b**), ruling out any effect of V_depol_ on the performance of the biosensor. In ST-mGlu5 expressing cells co-labeled with Lumi4-Tb and fluorescein (**Figure 3b**), quisqualate (10µM) induced a decrease in TR-FRET at V_rest_ (-17.2 ± 4.7%), reporting an agonist-induced increase of mGlu5 receptors in an active-like conformation. However, TR-FRET intensities measured in presence of quisqualate were significantly increased at V_depol_ (+7.0 ± 2.6%) compared to V_rest_, indicating that V_depol_ reduced the ability of the agonist to trigger conformational activation of the receptor (**Figure 3b**). This lower efficiency of quisqualate at V_depol_ to decrease the TR-FRET reports a higher proportion of mGlu5 receptors in an inactive-like conformation at V_depol_ compared to V_rest_. Consistently, we noticed a shift in IC_50_ of the LY341495 mGlu competitive antagonist, from 31.28 µM at V_rest_ to 6.75 µM at V_depol_ (**Figure 3c**), demonstrating a V_m_ effect on ligand affinity for the mGlu5 receptor orthosteric pocket. These results demonstrate the influence of V_m_ on the conformation of the receptor. V_depol_ stabilizes the inactive-like conformation of the receptor, increasing the antagonist potency.

**Figure 3.**
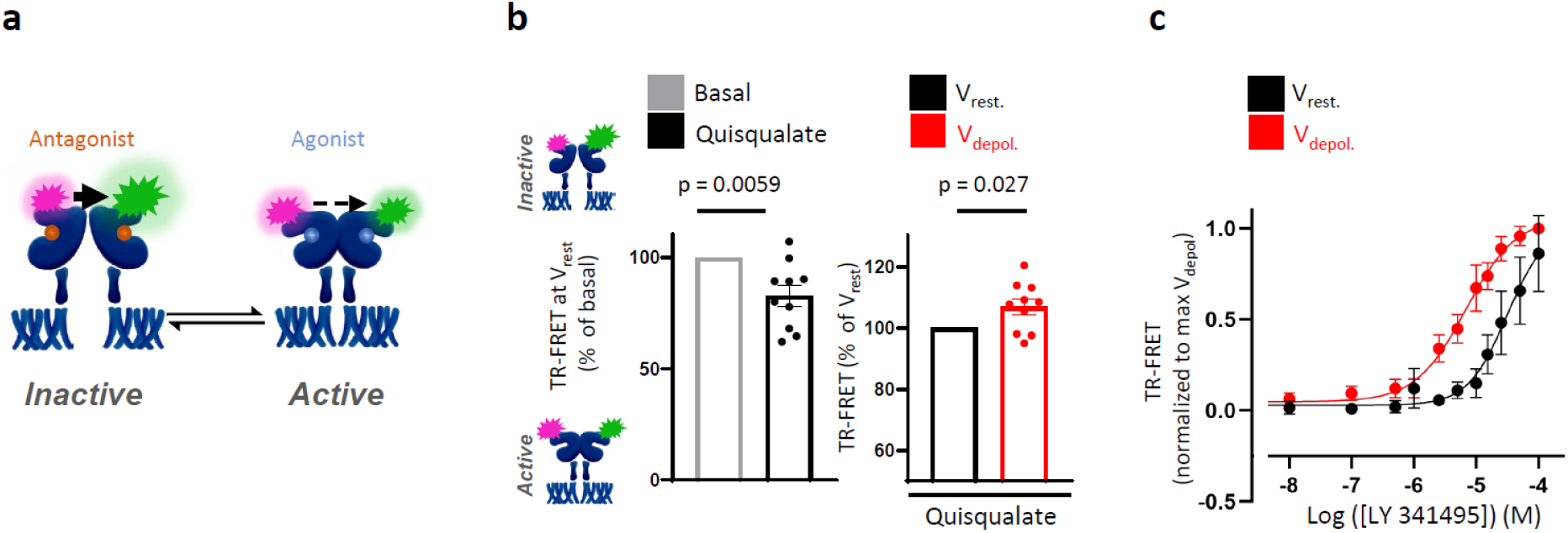
Voltage tunes mGlu5 receptor activation probability. **a)** Scheme of the TR-FRET sensor of mGlu5 conformation (38) with SNAP-Lumi4-Tb (purple) and SNAP-fluoroscein (green) linked on SNAP-tag-mGlu5 VFT domains. The FRET signal is inversely proportional to the number of receptors in an active-like conformation. **b)** TR-FRET intensity at V_rest_ with and without quisqualate 10µM (left) and quisqualate (10µM) effect at V_rest_ and V_depol_ (right). Data are mean ± SEM of n = 10 independent experiments. Statistics: Wilcoxon test. **c)** Dose-response curve of LY341495 mGlu antagonist applied at V_rest_ or V_depol_. Data are mean ± SEM of n = 4 independent experiments normalized to the maximum TR-FRET of V_depol_.

### V_m_ modulates mGlu5-mediated G_q/11_ activation

We then assessed the influence of V_m_ on the activation by mGlu5 receptor of canonical effectors, starting with the G_q/11_ activation. We monitored G_q/11_ activation by mGlu5 using Effector Membrane Translocation Assay (EMTA) ebBRET (40) in single-cell BRET imaging ((41), **Figure 4**) and cell population (**Figure Supp. 3**). HEK293T cells were transfected with mGlu5, G_q/11_, p63hRhoGEF-RlucII and rGFP-CAAX coding plasmids. G_q/11_ activation triggers the recruitment of its cytosolic effector, p63RhoGEF-RlucII, to the plasma membrane, which is reported by an increase in ebBRET with the plasma membrane-anchored rGFP-CAAX. The ebBRET signal is therefore a measure of G_q/11_ activation ((40), **Figure 4a**). Of note, the ebBRET signal was not affected by V_depol_ compared to V_rest_, validating the use of this biosensor to assess the influence of V_m_ on mGlu5-induced G_q/11_ activation (**Figure Supp 1c**). Application of DHPG (100µM) induced a slight but significant increase in ebBRET at V_rest_ and V_depol_ (**Figure 4b** and **4c**). Paired measurement of the DHPG net effect from individual cell ebBRET values revealed a mean DHPG-induced increase of 97 ± 9.9 milliBRET units at V_rest_, and only 22.8 ± 6.2 at V_depol_ (**Figure 4d**). These findings indicate a significant reduction of mGlu5 receptor-induced G_q/11_ activation at V_depol_. Similar results were obtained with ebBRET measurement in cell populations, where the mGlu5 agonist-induced increase of ebBRET was significantly reduced by V_depol_ (agonist-induced ebBRET increase was reduced by 24.8 ± 7.2 % at V_depol_ compared to Vrest, **Figure Supp 3a**). The same protocol applied to AT1 receptor showed no influence of V_m_ on AT1 receptor agonist-induced G_q/11_-coupling (**Figure Supp 3b**), suggesting once again that V_depol_ specifically affects mGlu5 receptor signaling.

**Figure 4.**
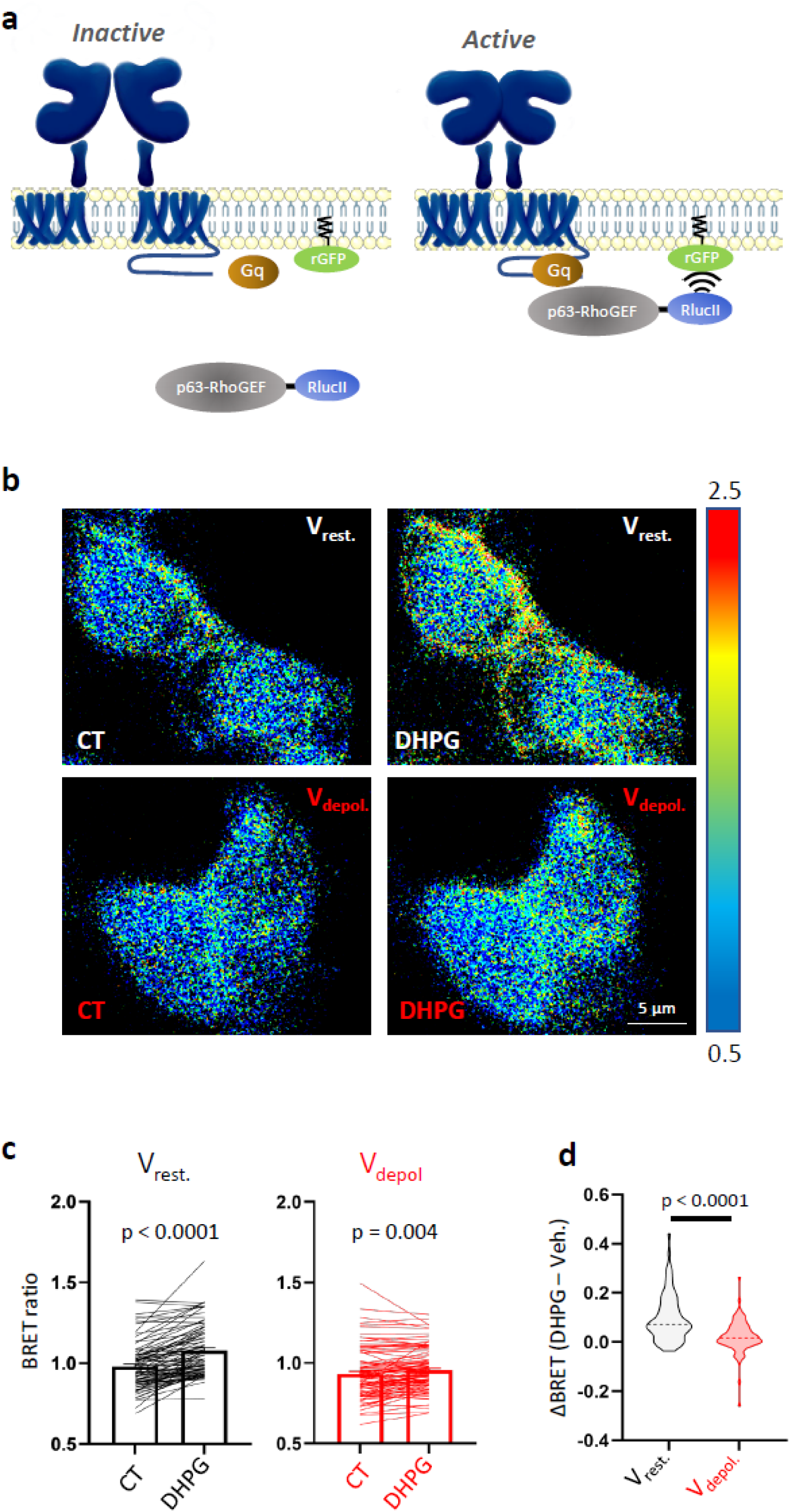
Voltage modulates mGlu5-mediated Gq activation. **a)** Scheme of EMTA ebBRET sensor of Gq activation (40). When Gq is activated by mGlu5, p63-RhoGEF-RlucII is recruited to the membrane where it interacts with rGFP-CAAX. **b** and **c)** Representative illustrations (b) and mean ± SEM (c) of BRET signals measured before (CT) and after (DHPG, 100µM) stimulation at V_rest_ or V_depol_. N= 86 to 97 cells from more than 3 independent experiments; statistics: Paired t-test. **d)** Single cell BRET measurement of DHPG net effect; statistics: Unpaired t-test.

### V_m_ affects mGlu5 gating of TRPC6 channels

Transient Receptor Potential Channels (TRPC) are well known effectors of GPCRs intracellular secondary messengers. Typically, TRPC6 opening is triggered by the PLC signaling cascade that triggers formation of diacylglycerol (DAG) and inositol 1,4,5-trisphosphate (IP3, (42)). In particular, mGlu5 receptor induces TRPC6-dependent Ca^2+^ influx (43,44). We then further tested the influence of V_m_ on the ability of mGlu5 to gate TRPC6 channels. For this purpose, we co-transfected mGlu5-Venus and TRPC6-tomato in HEK293T cells and recorded whole-cell currents induced by DHPG (100µM), in voltage-clamp experiments (**Figure 5)**. In cells co-expressing mGlu5 and TRPC6 (**Figure 5b**), but not in cells expressing mGlu5 alone (**Figure 5a**), DHPG applied at a holding potential of -80mV triggered a large inward current with similar kinetics and amplitude than previously reported in HEK293T cells (45). The current-voltage relationship (I/V curve, **Figure 5c**) also displayed typical TRPC6 permeation and rectification properties (46). Given that the ions flow through open channels is governed by the membrane potential, we could not simply compare the current density generated and recorded by activation of mGlu5 at different potentials. We therefore perfused the mGlu5 agonist at different holding potentials (either -80 mV or -20 mV) and then rapidly recorded the currents generated over the full range of V_m_ ramping (from -80mV to +60mV in 100ms, **Figure 5c**). When DHPG was applied at a holding potential of -20mV instead of -80 mV, the current amplitude recorded with the same voltage ramp protocol was strongly reduced. For example, we recorded a mean inward current density of 46.65 ± 10.32 pA/pF at -80 mV triggered by DHPG application at -80mV, versus 17.69 ± 6.36 pA/pF at -80 mV when DHPG was applied at - 20mV (**top insert, Figure 5c**). Importantly, these two protocols triggered the same I/V curve in non-stimulated cells, excluding a holding potential-dependent artefact during the recordings (**Figure Supp 4**). These results confirm that mGlu5 is more active at resting potentials compared to depolarized potentials, which impacts TRPC6 gating by mGlu5 downstream effectors.

**Figure 5.**
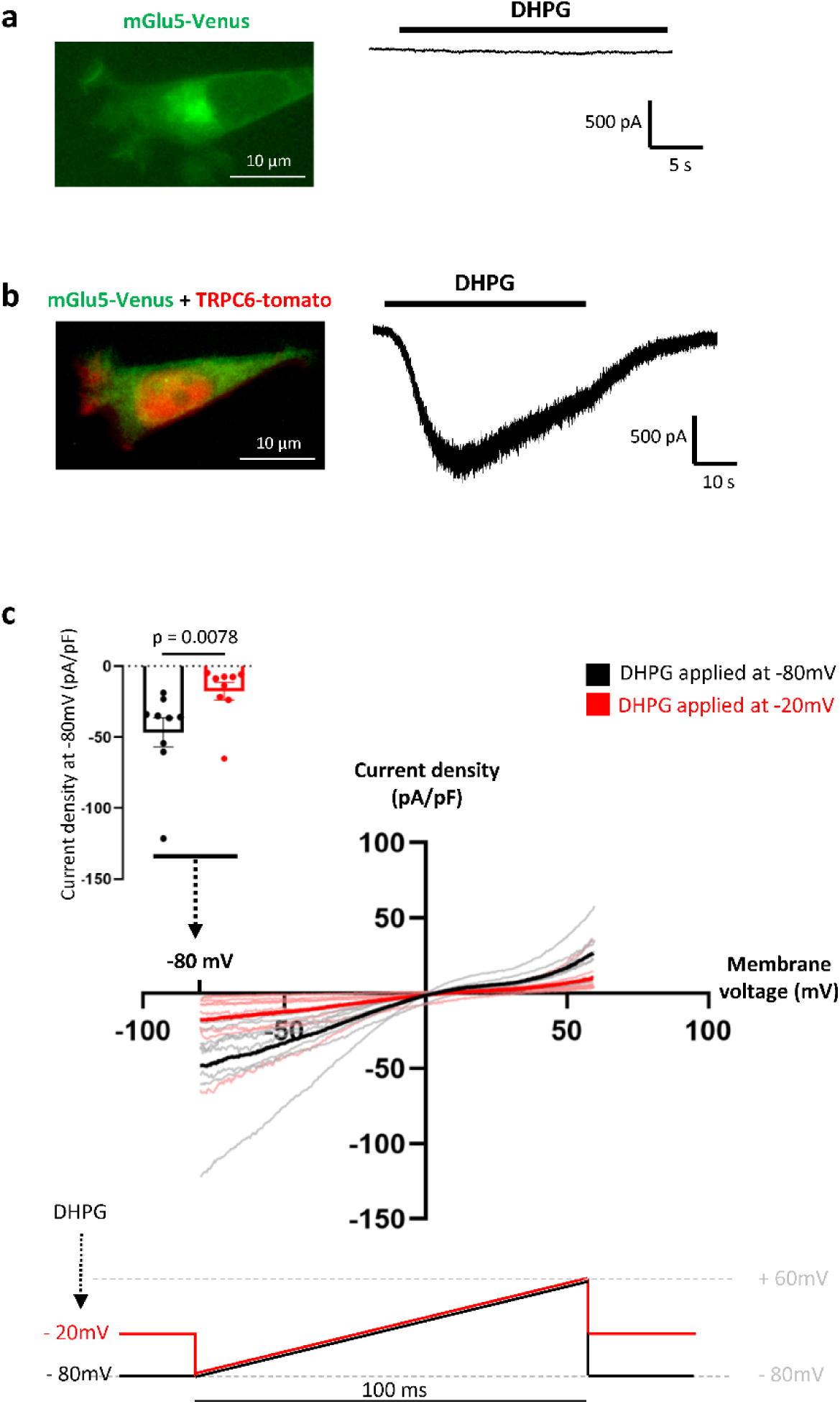
Voltage controls mGlu5 gating of TRPC6 channels. **a** and **b) -** Whole-cell patch clamp current induced by DHPG (100µM) on HEK293T cells expressing mGlu5-Venus alone (a) or with TRPC6-tomato (b). **c)** mGlu5-Venus/TRPC6-tomato co-transfected cells were held at -80mV (black) or -20mV (red) during DHPG (100µM) application. Current-voltage relationship was then recorded at the maximum response induced by DHPG (100µM) for potentials ranging from -80mV to +60mV in 100ms (bottom). Inset: Mean ± SEM current density at -80mV. Data are mean ± SEM of n = 9 cells per condition from 4 independent experiments, Statistics: Mann-Whitney test.

### V_m_ regulates mGlu5-NMDA receptors crosstalk in hippocampal neurons

The NMDA receptor conductance is well established to be regulated by type I mGlu receptors (47). Specifically, NMDA receptor potentiation by mGlu5 receptors involves the G_q_-protein-coupled receptor (GPCR) pathway, which includes protein kinase C (PKC) and Src signaling, in various neuronal contexts (24–26,48–50). In this study, we aimed to investigate whether V_m_ could regulate the crosstalk between mGlu5 and NMDA receptors in primary cell cultures of hippocampal neurons (**Figure 6**). To avoid stimulation of mGlu1 receptors, we used the mGlu1-specific negative allosteric modulator (NAM), CPCCOEt (100 µM), while stimulating mGlu5 receptors with DHPG. We recorded NMDA-induced currents in the absence of magnesium (Mg^2+^) to prevent voltage-dependent blockade of NMDA receptors. When applied alone at a holding potential of -80 mV, DHPG (50 µM) had a negligible effect (1.046 ± 0.45 pA/pF). In contrast, NMDA (30 µM) induced an inward current of 23.4 ± 3.7 pA/pF (**Figure 6a**), which was potentiated by 39.5 ± 6.5% by DHPG when applied at -80 mV (I_NMDA + DHPG_/I_NMDA_ ratio measured just before and after DHPG application, **Figure 6b, 6c left**). However, at a holding potential of -40 mV, DHPG only induced a 25.4 ± 5.2% potentiation of NMDA current density (**Figure 6b, 6c right**). Paired measurements of I_NMDA + DHPG_/I_NMDA_ performed subsequently at -80 mV or -40 mV on the same neuron in a random order confirmed a reduction of DHPG-induced NMDA receptor potentiation at -40 mV (+25 ± 4.8%) compared to -80 mV (+38.11 ± 8.2%) (**Figure 6d**). These results demonstrate the crucial role of V_m_ in the control of mGlu5-NMDA crosstalk in neurons, and they corroborate our previous findings on the global voltage-dependence of mGlu5 receptor activity, which is inhibited by depolarized membrane potentials.

**Figure 6.**
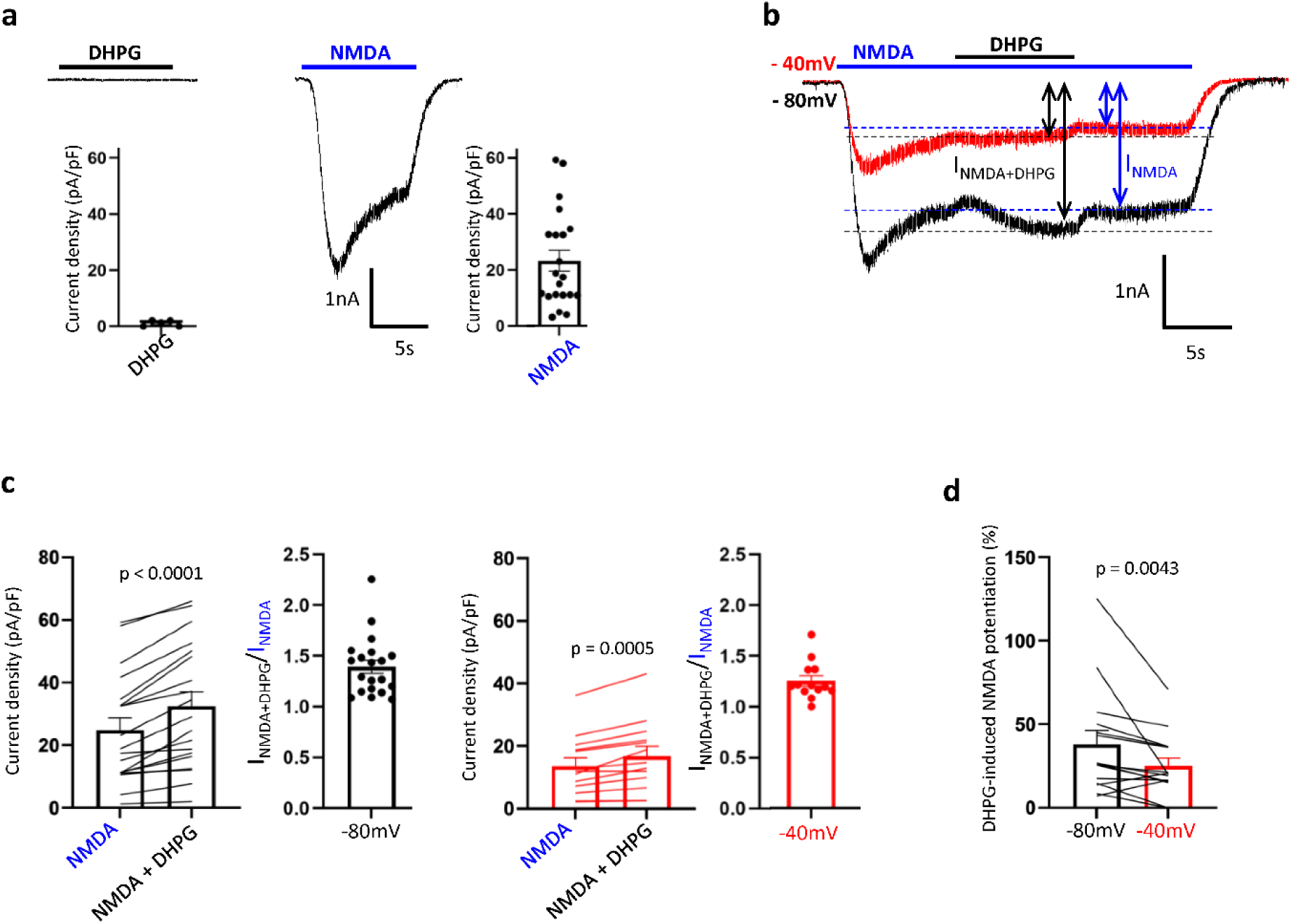
Voltage regulates mGlu5 gating of NMDA receptor in hippocampal neurons. **a)** Representative current and mean density induced by DHPG (50µM, left) or NMDA (30µM, right) at -80mV. **b - d)** Representative currents (b), mean current density and I_NMDA+DHPG_/I_NMDA_ current ratio (c), and percentage of DHPG-induced NMDA current potentiation (d) measured in presence of NMDA (30µM) alone or co-applied with DHPG (50µM) at -80mV (black) and - 40mV (red) holding potential. Measures of I_NMDA_ (blue) and I_NMDA+DHPG_ (black) used for quantification in c and d are indicated by arrows in b. In d, paired measurement of I_NMDA + DHPG_ / I_NMDA_ were performed subsequently at -80mV or -40mV, in a random order, on the same neuron. c and d data are mean ± SEM of n = 10 to 15 neurons from at least 4 independent experiments; Statistics: Wilcoxon matched-pairs signed rank test.

Therefore, the optimal functioning of the mGlu5 receptor, which enhances NMDA receptor activity in neurons, occurs at the resting potential of neurons. This finding appears to conflict with the fact that NMDA receptors are typically blocked by Mg^2+^ at resting membrane potential. Activation of the NMDA receptor indeed requires depolarization of the postsynaptic element to release magnesium from the channel pore, and simultaneous binding of glutamate released from the depolarized axon terminal. However, recent studies have shown that the physiological concentration of Mg^2+^ in the interstitial medium (0.7 mM, (51)) is lower than what is typically used experimentally in the ACSF composition to record ex vivo neuronal activity, suggesting that the NMDA receptor activity at resting potential in physiological concentrations of Mg^2+^ may have been underestimated (52). Furthermore, GluN2D-containing receptors may exhibit reduced Mg^2+^ sensitivity compared to GluN2A- or GluN2B-containing receptors (53). Therefore, we investigated the mGlu5-induced potentiation of NMDA receptors at resting potential in the presence of 0.7 mM Mg^2+^, by recording currents and calcium transients (**Figure 7**). Whole-cell current recordings revealed a residual NMDA-induced current of 1.80 ± 0.30 pA/pF density, which was potentiated to 2.40 ± 0.40 pA/pF by DHPG co-application (**Figure 7a**). The full current potential relationships are displayed in **Figure 7b**.

**Figure 7.**
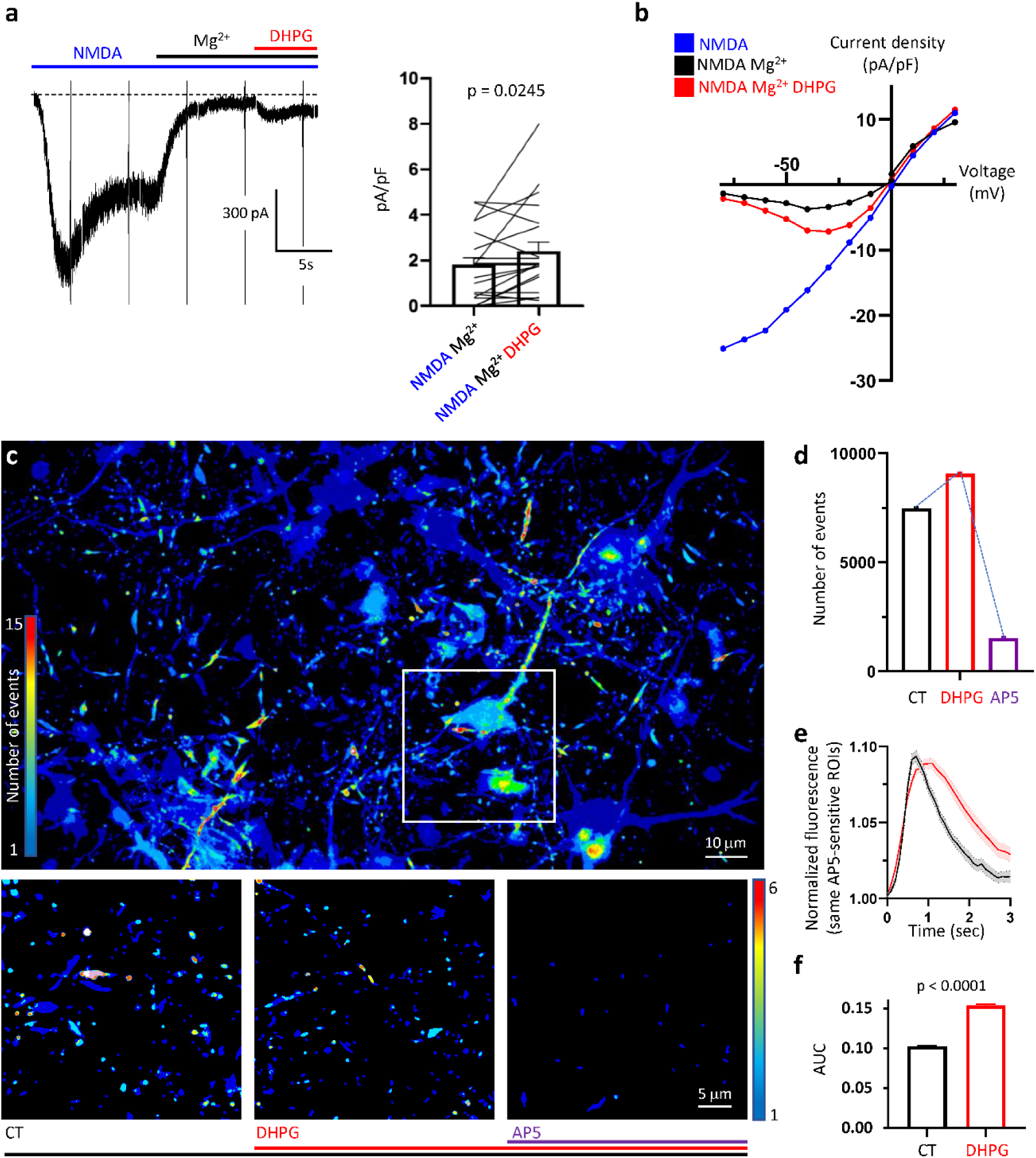
DHPG-induced potentiation of NMDA currents at resting potential, in physiological concentration of Mg^2+^. **a)** Left - Representative NMDA (100µM)-induced current, without (NMDA) or with (Mg^2+^) extracellular Mg^2+^ (0.7 mM), co-applied with DHPG (50 µM). Right – Mean ± SEM current density recorded in 0.7 mM Mg^2+^, from n = 21 neurons in 3 independent experiments. Wilcoxon matched-pairs signed rank test. **b)** Current density-voltage relationship (-80 -> +60 mV within 100ms every 5s) performed as illustrated in (a) during NMDA application (at the plateau response) without (blue) or with 0.7 mM Mg^2+^ (black) co-applied with DHPG (red). Mean from n = 8-10 neurons per condition. **c)** Projection of spontaneous calcium transients, recorded during 2 min and 15 sec, in Mg^2+^ (0.7mM) at resting potential of neurons in a representative field of view. Top: all events for the entire field of view. Bottom: small non-propagating calcium transients from the white square in the top image, for three consecutive acquisitions. Left: basal. Middle: basal + DHPG (50 µM). Right: basal + DHPG (50 µM) + AP5 (50 µM). **d)** Number of calcium transients recorded in the whole field of view before (CT) and after sequential application of DHPG and AP5. **e)** GCaMP6s fluorescence fluctuation during small non-propagating calcium transients, Mean ± SEM of 91 and 94 traces, before (black) and after (red) DHPG application, measured on 10, identical, AP5-sensitive ROI; **f)** Mean area under curve (AUC) of calcium transients. Each bar of the histogram is the Mean ± SEM of all AUC traces recorded in the whole field of view before and after DHPG application for 2 min and 15 seconds in both conditions. Statistics: Unpaired t test.

In a second set of experiments, we measured calcium fluctuations, still in presence of physiological concentrations of Mg^2+^ and following successive application of DHPG and AP5 (**Suppl Video 3**). GCaMP6s fluorescence fluctuations highlighted basal spontaneous Ca^2+^ transients, of various shape, size, and kinetics, as illustrated by a projection of all spontaneous Ca^2+^ transients recorded during 2 min and 15 sec (**Figure 7c**). The variety of these Ca^2+^ events certainly relies on the nature of the receptors and channels involved in their triggering. To focus on synaptic events, we selected small and non-propagating calcium transients. The majority of these events were blocked by AP5 (50 µM) at the end of the experiment, revealing NMDA-dependent events (**Figure 7d**). We could *a posterior* focused in these regions of interest to measure the influence of mGlu5 activation on NMDA function (**Figure 7e**). DHPG increased the number of basal calcium transients (**Figure 7d**) and their area under curve (AUC indeed significantly increased from 0.102 ± 0.001 to 0.154 ± 0.002 following DHPG application, **Figure 7f**). More importantly, DHPG changed the shape of the NMDA-dependent events inducing a more sustained calcium inflow over time (**Figure 7e**). Taken together, our data show that the mGlu5 receptor potentiates NMDA receptor activity at resting membrane potential, increasing currents and particularly calcium influx, thus expanding the functional importance of NMDA receptors to resting neurons.

## Discussion

In this study, we demonstrate that the activity and signaling of the mGlu5 receptor is influenced by the membrane potential of the cell, highlighting an aspect of the receptor that has previously been overlooked. Specifically, a depolarization of the membrane promotes an inactive-like conformation of the receptor, a reduction in mGlu5 coupling to G_q/11_ and subsequent release of Ca^2+^ from intracellular stores. This results in a reduced activation of downstream effectors, including TRPC6 channels and ionotropic NMDA receptors. The mGlu5 receptor is a critical component in the modulation of synaptic transmission, and it plays a role in both inducing and regulating Hebbian and homeostatic synaptic plasticity processes. As such, the sensitivity of the mGlu5 receptor to membrane potential will impact its functions, and its role may vary depending on the context of neuronal activity.

### The mGlu5 receptor functions optimally at the resting potential of cells

We observed that each of the receptor signaling steps we studied were affected by the membrane potential, and our results indicate that the optimal function of the receptor occurs when the cell is at rest.

A sensor that monitored the distance between the extracellular domains of the receptor’s protomers within the mGlu5 homodimer over time allowed us to determine the activation state of the receptor when bound to an agonist. FRET measurements showed that depolarization of the membrane favored an inactive-like conformation of the mGlu5 receptor, which increased the potency of its orthosteric competitive antagonist. These findings were consistent with a previous study demonstrating that glutamate binding to the mGlu3 receptor could be inhibited by membrane depolarization (11). Although this sensor provided information only about the extracellular domains of the receptor, this is a reliable indicator of the activation state of mGlu receptors (38,39). The conformational change of the extracellular domains of the receptor may be induced by a conformational change of the seven transmembrane domains (7TM) due to depolarization, as the 7TM are the closest elements of the receptor to the electric field (see discussion below). This hypothesis is supported by structural studies showing that activation of the 7TM stabilize the active conformation of the extracellular domains of the mGlu5 receptor (54).

The mGlu5 receptor’s ability to activate G_q_ is diminished under depolarizing conditions. The G_q_ protein activation sensor relies on the translocation of G_q_ protein effector, p63RhoGEF, to the membrane upon activation of a GPCR. The BRET signal observed between p63RhoGEF bound to RlucII and membrane-targeted rGFP through a CAAX sequence corresponds well with G_q_ protein activation (40). The translocation of the G_q_ effector to the membrane appears to be voltage-dependent only when the mGlu5 receptor is activated. In the exact same experimental conditions, no effect was observed on activation of the AT1 receptor. Nevertheless, this receptor activates the same pathway as the mGlu5 receptor (55–57), and therefore serves as a rigorous control to emphasize the specificity of V_m_ action on mGlu5 receptor signaling. Activation of G_q_ proteins by mGlu5 receptors induces IP3 production and triggers a significant release of Ca^2+^ from intracellular stores into the cytosol, resulting in typical Ca^2+^ oscillations previously reported to be caused by PKC-induced phosphorylation and desensitization of mGlu5 receptor (32). Depolarization reduced the release of Ca^2+^ from intracellular stores generated by mGlu5 receptor activation. Not only did the number of cells generating Ca^2+^ oscillations decrease following mGlu5 receptor stimulation, but also the frequency of oscillations per oscillating cell was significantly reduced by membrane depolarization. In contrast, V_m_ did not influence the Ca^2+^ response initiated by activation of the AT1 receptor, which is also known to produce similar Ca^2+^ release (57). These findings indicate that depolarization specifically impairs Ca^2+^ release induced by the mGlu5 receptor.

Activation of G_q_-protein by mGlu5 receptors induces production of PIP2 and DAG, which in turn triggers the opening of TRPC channels (42,45). Co-expression of TRPC6 channels in cells resulted in the inward current being triggered by the activation of mGlu5 receptors, with typical current kinetics, amplitude, and rectification properties (43,44), indicating that these channels are gated by mGlu5 receptor activation. By varying the imposed potentials for a few milliseconds, current-potential curves were used to determine the channel conductance. Stimulation of mGlu5 receptors at a holding potential of -20 mV instead of -80 mV significantly decreased the conductance of TRPC6. These findings suggest that membrane depolarization inhibits the ability of the mGlu5 receptor to open TRPC6 channels.

Other channels are controlled by the mGlu5 receptor, including the ionotropic glutamate receptors of the NMDA type (47), which also play a fundamental role in the induction of synaptic plasticity. The intricate cross-talk between mGlu5 and NMDA receptors defines functional neuronal networks, which evolve through complex mutual regulations that are influenced by the dynamics of synaptic protein complexes and the context of neuronal activity (25,58,59). These regulations can lead to either potentiation or inhibition of their activations. Potentiation of NMDA receptor activity by mGlu5 receptors has been shown to depend on the G_q_-PKC-Src signaling pathway in various neuronal contexts (24,25,48,49). Our experimental results confirm this finding, as we observed a significant increase in NMDA current upon mGlu5 stimulation. However, we also found that this facilitation is dependent on the holding potential of the neuron and decreases with depolarizing membrane potential. These results suggest that the activity of mGlu5 receptors is lower at depolarized potentials, limiting the facilitation of NMDA to potentials close to the resting potential of neurons. In addition, our findings reveal a permissive role of the mGlu5 receptor on NMDA receptor activation even under physiological concentrations of Mg^2+^ at resting potential, further expanding the role of NMDA receptors in synaptic plasticity.

### Reconsideration of mGlu5 and NMDA functions

Our study highlights the crucial role of membrane potential in modulating the activity of mGlu5 receptors, which are involved in synaptic transmission, plasticity, and metaplasticity. We found that mGlu5 receptors are fully active at resting membrane potentials, but their activity is dampened upon depolarization. This sensitivity to V_m_ is highly relevant given that neuronal activity can cause fluctuations in V_m_. Additionally, our study suggests that the optimal functioning of mGlu5 receptors at resting membrane potential may potentiate NMDA receptors too. Traditionally, NMDA receptors are known to be “coincidence detectors” of pre- and post-synaptic activation because they are blocked by the Mg^2+^ ion at resting potential (22,23). However, our results indicate that they may also play an unexpected role at resting potential, facilitated by the permissive effect of mGlu5 receptors. Recent measurements of interstitial fluid composition in vivo show that the actual Mg^2+^ concentrations (0.7 – 0.8 mM. (51,60)) are lower than those used experimentally, which may underestimate the functional importance of NMDA receptors at resting potential (52). Our findings add complexity to the functional cross-talk between mGlu5 and NMDA receptors, with mGlu5 acting as a starter for NMDA receptor activation on recently resting neurons. Once the membrane potential is depolarized enough to unblock NMDA receptors, they can function autonomously, while negative feedback from V_m_ on mGlu5 receptors limits NMDA facilitation and synapse overexcitability. This finding is likely to have significant implications for the nature of induced neuronal plasticity, which relies on the concerted activities of mGlu5 and NMDA receptors (61,62). Different cellular mechanisms for inducing plasticity will come into play depending on the type of activatable receptor following network activity.

The acute effect of depolarization on the activity and signaling of mGlu5 receptors adds to the extensive regulation of protein complexes around these receptors and their functional interactions that are regulated by neuronal activity. The mGlu5 receptor has been proposed as a homeostatic regulator of synaptic transmission, a role that requires activity-induced monomeric Homer1a expression (27,58,63). Within the neuronal environment, the function of mGlu5 receptors is dynamically regulated by interactions with multimeric Homer proteins, which is dampened by the monomeric Homer1a protein during sustained synaptic activity (64,65). Our team recently demonstrated an instantaneous disruption of the mGlu5-Homer interaction following membrane depolarization in hippocampal neurons (34), which cannot be explained by the induction of monomeric Homer1a synthesis, a process taking place several minutes later. This depolarization-dependent disruption occurs even if NMDA receptors are blocked (34) and may be attributed to the direct sensitivity of the mGlu5 receptor to membrane potential, as demonstrated in this study. However, testing this hypothesis would require knowledge of and mutations to the voltage sensor of the mGlu5 receptor without affecting the receptor’s response to its ligand.

Following the induction of a first plasticity event, the expression of Homer1a blocks the activity of the mGlu5 and NMDA receptors to prevent the induction of subsequent plasticity, allowing cellular signaling required for the expression of the engaged plasticity and the maintenance of the neuron in the functional network in which it has been committed (34). It is worth noting that the direct effect of V_depol_ on mGlu5 receptor functioning described in this study is similar to the effect of Homer1a, which inhibits canonical receptor signaling. The acute effect of depolarization is likely to control the function of the mGlu5 receptor on a shorter time scale. The decreased affinity of the receptor for multimeric Homer, first by membrane depolarization (seconds) and then by CamKII-dependent Homer phosphorylation (minutes, (66)), would promote decreased competition and enable the interaction with monomeric Homer1a once this protein is expressed (20-30 minutes, (65)), enabling long-lasting plasticity.

### The mGlu5 receptor senses moderate V_m_ variations, similar to natural fluctuations of membrane potential occurring at synapses

In this study, we have demonstrated that a modest increase in membrane potential by approximately 20-25 mV is sufficient to reduce mGlu5 receptor activity by half. Our findings reveal that the mGlu5 receptor is highly sensitive to changes in membrane potential compared to other GPCRs which require stronger depolarizations, at least twice the level used in our study, and sometimes up to 140 mV, to induce a similar shift in receptor efficacy (4,67,68). Both patch clamp and KCl depolarization methods resulted in the same polarity of membrane potential effect on mGlu5 receptor function. Moreover, the effect of depolarization was comparable when applied before or during agonist perfusion, suggesting that membrane potential can either prevent optimal activation or rapidly reverse ongoing activation of the mGlu5 receptor. Notably, changes in dendritic spine membrane potential during post-synaptic potentials have been reported to be in the range of 20-25 mV, a magnitude similar to that observed in our study (69–71). Thus, local synaptic potentials may mediate rapid modulation of mGlu5 function in neurons. Although membrane potential changes were applied tonically in our study, similar results have been obtained by others using physiological frequency ranges, mimicking trains of action potentials (12,15). A more resolutive investigation would provide better insights into the intensity and duration of membrane potential variations required to modulate mGlu5 receptor activity and refine our understanding of the type of neuronal activity of similar magnitude and temporal order that could affect mGlu5 function.

### Membrane depolarization controls mGlu5 receptor activity, as a NAM

The scope of this study did not include the identification of the voltage sensor within the mGlu5 receptor. However, previous studies on other GPCRs have identified molecular determinants of V_m_ sensitivity, such as the binding sites of orthosteric or allosteric ligands, G protein binding domains, or Na^+^ ions within the 7TM (72). These studies mainly concern class A GPCRs, where the molecular sites mentioned above are located in the 7TM. Unfortunately, very little is known about the voltage-sensing domains of class C GPCRs, to which the mGlu5 receptor belongs. Nevertheless, it is reasonable to assume that the voltage sensor is also located in the 7TM, directly exposed to the magnetic field. In the atypical structure of class C GPCRs, the extracellular domains of the receptor (and the orthosteric binding site) are approximately 10 nm distant from the membrane (54), out of the membrane electric field, which excludes their involvement as a voltage sensor. In contrast, allosteric modulators bind to the 7TM of the receptor. Interestingly, many studies show that V_m_ affects the efficiency or agonist potency of class A GPCRs in a manner similar to allosteric modulators (12,67,68). Specifically, tyrosine residues located in the allosteric site are involved in V_m_ detection by GPCRs (10). Of note, the allosteric binding site of the mGlu5 receptor also contains a tyrosine residue (Tyr6593.44) that enables NAM inhibitory effects (54,73).

Functional data collected in this study suggests that membrane depolarization causes a NAM effect on the mGlu5 receptor. First, NAMs cause a decrease in agonist efficacy with virtually no changes on agonist potency (18), similar to the effect of depolarization on the dose-response curve (**Figure 1e**). Second, mGlu5 receptor-mediated Ca^2+^ oscillations are known to undergo frequency reduction by NAMs, such as MPEP, supported by a decrease in the proportion of oscillating cells (35,36). Interestingly, a similar process is induced by membrane depolarization, namely, a reduction in frequency when depolarization is applied before or after the initiation of oscillations, correlated with a decrease in the proportion of oscillating cells (**Figure 2**). Finally, the competitive antagonist LY341495 binds to the Venus Flytrap (VFT) domains, and its binding is favored by membrane depolarization, acting like a NAM that stabilizes the inactive conformation of the receptor. Similarly, mGlu4 and mGlu5 allosteric modulators binding to the 7-TM of the mGlu receptor can drive changes in VFT domain conformation (54,74), meaning that 7-TM modulation by V_m_ could, as a NAM, be transduced on VFT domain conformation. Hence, membrane depolarization would reduce the activity of receptors expressed at the membrane, like a NAM. These results support to search for the voltage sensor of mGlu5 in the NAM binding site.

In conclusion, this study has identified mGlu5 receptor as a member of the small yet expanding group of voltage-sensitive GPCRs. The high sensitivity of mGlu5 receptor to changes in membrane potential at synapses suggests that its effects extend beyond canonical signaling and may even impact its cognate NMDA receptor. Optimal functioning of mGlu5 receptor at hyperpolarized potentials could have far-reaching consequences, such as restricting its activation to specific locations by integrating glutamate spill-over and electrical propagation at synapses. Additionally, mGlu5 receptor may exert cell-type-dependent actions, conditioned by the resting potential of cells. Finally, given the crucial role of this receptor in neuronal physiology, it is likely that its voltage sensitivity plays a critical role in pathological conditions associated with variations in intrinsic excitability.

## Methods

### Plasmids

Plasmids were amplified using E.Coli DH5α strain (Life Technologies). DNA was then purified using the NucleoBond^®^ Xtra Plus Midi-Prep kit (Machery-Nagel^®^). The plasmids used were: pcDNA3.1-rGFP-CAAX (37), pcDNA3.1-p63RhoGEF-RlucII (40), pcDNA3.1-Gαq (gift from Jean-Philippe Pin), pRK5-SNAPTag-AT1 (CIS bio international), pRK5-Myc-mGlu5a and pcDNA3.1-Myc-mGlu5a-Venus (75). pCAGGS-TRPC6-ires2-mbtdtomato was obtained by cloning mouse TRPC6 sequence in pCAGGS plasmid (76) upstream the sequences of IRES2 and membrane-anchored tdtomato.

### Reagents

(S)-3,5-Dihydroxyphenylglycine (DHPG, HelloBio^®^); Quisqualate (Tocris); LY341495 (Tocris^®^); 2-methyl-6-(phenylethynyl)-pyridine hydrochloride (MPEP, HelloBio^®^); human angiotensin II (HelloBio^®^); N-Methyl-D-Aspartate (NMDA, Tocris^®^). 7-(Hydroxyimino)cyclopropa[*b*]chromen-1a-carboxylate ethyl ester (CPCCOEt, Tocris^®^); Tetrodotoxin (TTX) Citrate (Latoxan^®^), Bicuculline (Hellobio^®^). Except for dose response curves, all the mGlu5, AT1 and mGlu1 ligands are applied at saturating concentrations.

### Cell cultures and transfection

HEK293T cell lines grown in DMEM GlutaMAX^®^ medium (Gibco^TM^) supplemented with 10% fetal bovine serum and 1% penicillin/streptomycin were maintained at 37°C in a humidity-controlled atmosphere with 5% CO_2_. The ST-hmGlu5 HEK293T cell line enables inducible expression of the hmGlu5 receptor containing a SNAP-tagged extracellular VFT domain, as previously described (77). Briefly, the release of the constitutive Tet repression of the ST-hmGlu5 gene is induced by doxycycline (Dox), allowing a reproducible and controlled expression, proportional to Dox concentration and incubation time. Cells were maintained in ST-hmGlu5 transgene selection medium in the presence of 100 μg/ml hygromycin B and 15 μg/ml blasticidin. Dox (1µg/ml) was added 6h (Ca^2+^ experiments) or 10 to 24h (TR-FRET measurement) before experiments. Cell lines transduction with the pWPT-EF1a-GcaMP6s-P2A-Scarlet lentivirus (gift from Vincent Compan) enabled stable expression of GcaMP6s fluorescent Ca^2+^ sensor. Plasmids transfection was performed on adherent cells the day after the seeding in antibiotic-free culture medium using Lipofectamine 2000^TM^ (Life Technologies) diluted in OptiMEM GlutaMAX^®^ (Life Technologies) or jetPEI^®^ (Polyplus, only for patch clamp experiments), 24 to 48h before experiments.

Primary cultures of hippocampal neurons were prepared from C57BL/6 mice P0-2 pups as previously described (78). Neurons were seeded onto plastic dishes coated with Poly-L-ornithine (0.03 mg/ml) and laminin (1µg/ml) in Neurobasal, B27, Glutamax, Glutamine, serum and 1% penicillin/streptomycin medium. Neurons were then maintained at 37°C with 5% CO2 and incubated from DIV2 to DIV3 with AraC (1 µM) to inhibit glial cell proliferation. The medium was then progressively replaced by a serum-free medium (BrainPhys, B27, Glutamax).

### Recording Solutions

In all experiments, the extracellular solution contained (in mM): NaCl 140, CaCl_2_ 2, MgCl_2_ 0 to 2, KCl 3, HEPES 10, D-glucose 10 (pH = 7.4, 330 mOsm). For neurons, Glycine 0.01, bicuculline 0.01, and tetrodotoxin 0.0003 were added the day of the experiment. For whole-cell voltage clamp recordings, the pipette was filled with an intracellular solution (in mM): CsCl 140, EGTA 5, CaCl_2_ 0.5, HEPES 10, ATP-Na_2_ 2 and D-glucose 10 (pH = 7.2, 300 mOsm). Equimolar solutions containing increased KCl concentrations (from 3mM to 100 mM) were obtained by equal decrease of NaCl concentration.

### Ca^2+^ measurement

- Cell population: live Ca^2+^ fluctuations were recorded at 37°C on ST-hmGlu5 GcaMP6s-P2A-Scarlet HEK293T cells, or when specified on GcaMP6s-P2A-Scarlet HEK293T cells transfected with pRK-SNAPTag-AT1 (25 ng/well), seeded at 5 x 10^4^ cells per well in 96-wells plates with transparent bottom, with the FDSS/µCell plate reader (Hamamatsu^®^) – Acquisitions of 1Hz; Exc: 480 nm – Em: 540 nm; high-speed digital EM CCD camera. Each independent experiment was performed in triplicate, which average corresponds to n = 1. Data were normalized to the maximum response at V_rest_ in each independent experiment. Basic calculations from the raw data files were performed on Microsoft Excel (Office 2016), statistical analyses and graphics on GraphPad Prism 8.1.
- Single cell imaging: GcaMP6s fluorescence fluctuations were recorded on ST-hmGlu5 GcaMP6s-P2A-Scarlet HEK293T cells, using an Axio Observer 7 KMAT fluorescence microscope (Carl Zeiss), equipped with a Plan-Neofluar 40x/1.30 EC oil objective (M27, ZEISS®), Exc 470/40 nm – Em 525/50 nm filters and ORCA-Quest qCMOS camera (Hamamatsu); all controlled with Metamorph software. Images on HEK cells were acquired at 3Hz and analysed with Fiji using a custom code computing the average fluorescence intensity of the regions of interest. For the frequency analysis, MATLAB was used to automatically detect the oscillations occurrences and compute the instantaneous and global oscillation frequencies. First, baselines were computed as the 10^th^ percentile of the trace using a sliding window of size 400 frames. Signal was then divided by the baseline. Oscillations were detected using the function findpeaks with a minimal peak prominence of 0.01. Instantaneous frequency was computed as the number of oscillations minus one divided by the time between first and last oscillation. Global frequency was computed as the number of oscillations divided by the total duration. Spontaneous Ca^2+^ transients in neurons infected with the pWPT-CAMKIIa-GcaMP6s-P2A-Scarlet lentivirus were acquired at 10 Hz, for series of 2 min and 15 sec. Image stacks were then analyzed as follow: images were filtered with a 5*5 median kernel. Each pixel was divided by its average value over the whole stack. Each pixel was gaussian filtered along the time axis with a sigma of 2 frames. Time derivative was applied to the stack with a step of 6 frames. Images were thresholded with a value of image mean + 6 * image standard deviation. A filter was then applied to keep only voxels with at least 10 thresholded voxels in their 3*3*3 neighborhood, to remove isolated voxels. Thresholded voxels spatially connected were assembled as events. Maximal time projection of those events gave their spatial footprint: the list of pixels in the image. Events with lower than 16 pixels were rejected. Events were classified in different categories based on two criterions: a threshold of 1000 pixels separating big events from small events, a threshold of 0.8 px of propagation distance separating events propagating or not. Propagation distance was measured as the difference of position between the centroid of the pixels thresholded at the beginning and at the end of the onset. Both thresholds were applied to focus on “small, non-propagating calcium transients” at synapses. Projection images were obtained by accumulation of the events footprint. Fluorescence time traces were extracted for each event as the average fluorescence over the event pixels and normalized by the fluorescence just before the onset of the event. Area under curve was computed between the onset and 4 seconds later.

### TR-FRET measurement

TR-FRET measurements were performed as previously described (39). Briefly, ST-hmGlu5 HEK293T seeded at 5 x 10^4^ cells per well in 96-wells black bottom plates were incubated for 1h-1h30 with 100nM of SNAP–Lumi4-Tb and/or 60 nM of SNAP–green. Measurements were performed at room temperature using the Pherastar FS (BMG LabTech). Following SNAP–Lumi4-Tb excitation (337 nm), the decays of SNAP–green or SNAP–Lumi4-Tb emission were measured every 10µs at 520 nm and 620 nm respectively. TR-FRET ratios and SNAP–Lumi4-Tb decays were calculated following Scholler et al. recommendation for mGlu receptors with the corresponding time intervals: 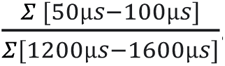. For cells incubated only with SNAP–green, the fluorescence was measured at 520 nm with a 485 nm excitation. Each independent experiment was performed in triplicate, which average corresponds to n = 1 and data were normalized to the maximum response at V_rest_ except for LY341495 dose-response curves which were normalized to the maximum response at V_depol_. Basic calculations from the raw data files were performed on Microsoft Excel (Office 2016), statistical analyses and graphics on GraphPad Prism 8.1.

### BRET measurement

After coelenterazine H (Coel H, Promega, 2.5 µM) addition, ebBRET measurements in cell population were performed using the LB940Mithras plate reader (Berthold) as previously described (37,40), on cells seeded at 5 x 10^4^ cells per well in 96-wells white bottom plates and transfected with pcDNA3.1-rGFP-CAAX (33,3 ng/well), pcDNA3.1-p63RhoGEF-RlucII (5,1 ng/well), pcDNA3.1-Gαq (6,75 ng/well) combined with the pRK5-Myc-mGlu5a (14,7 ng/well) or pRK-SNAPtag-AT1 (14,8 ng/well). BRET, or the 535 nm/485 nm ratio, was assessed by calculating the ratio of the light emitted by the acceptor (510–550 nm band-pass filter, Em535) to the light emitted by the donor (460–500 nm band-pass filter, Em480).

Single cell BRET imaging was previously described (41). Acquisitions were performed on cells seeded in 35mm diameter glass dishes (MatTek® for microscopy) coated with poly-ornithine (0.03 mg/ml, for 30min), after Coel H (5 µM) addition, using the Axio Observer 7 KMAT fluorescence microscope (Carl Zeiss), equipped with a Neofluar 40X/1.3 planar EC oil objective (M27, ZEISS) and an ORCA-Quest qCMOS camera (Hamamatsu), controlled by Metamorph software. The emission of RlucII (10s exposure, FF01-450nm/70-25 Brightline filter, Semrock) and rGFP (20s exposure, HQ535nm/50M filter, Chroma Technology Corp) were measured successively. Images were analyzed using the BRET Analyzer macro on Fiji (79). Overall, this analysis uses an open-source toolkit for Fiji (https://github.com/ychastagnier/BRET-Analyzer) performing four key steps in the analysis: (1) background subtraction from the image, (2) image alignment over time, (3) composite thresholding of the image, and (4) pixel-by-pixel division of the image and distribution of the BRET ratio intensity on a pseudo-color scale. The BRET ratio corresponds to the acceptor/donor emission ratio (535/450).

### Electrophysiology

HEK293T cells, seeded in 35 mm diameter plastic dishes at low confluency (1×10^4^ cells) and transfected with pcDNA3.1-Myc-mGlu5-Venus (0.25 µg) and pCAGGS-TRPC6-ires2-mbtdtomato (0.5 µg) plasmids, were targeted by fluorescence microscopy. Measurements were performed in the whole cell configuration, in voltage or current clamp mode, with pipettes of 2-4 MΩ resistance when filled with the appropriate intracellular medium.

At 10 to 12 days in vitro, hippocampal neurons were used for endogenous NMDA current measurements. Patch clamp recordings were performed in the whole-cell configuration with pipettes of 3-5 MΩ resistance when filled with the appropriate intracellular medium. NMDA stimulations were separated by 2 min to avoid receptor desensitization. All recordings were performed at room temperature for less than 1 h per dish. Drugs were applied using a gravity perfusion system allowing complete exchange of the cellular environment in less than 30ms.

Currents were recorded using an Axopatch 200B amplifier (Axon Instruments), with noise removal at 50-60 Hz (Hum Bug Noise, Quest Scientific) and digitized at 3 kHz before being stored in Clampex software (version 8.1). The data were then analyzed using the Clampfit 11.2 software from Axon instruments (Molecular Devices).

## Acknowledgments

The authors thank the iExplore animal facility (IGF, Montpellier) and the Arpege platform (IGF, Montpellier) for the use of plate reader fluorimeter for cell-population BRET and FRET assay. The authors thank Philippe Marin, Jean-Philippe Pin, Laurent Prézeau, Charlotte Avet and Michel Bouvier for providing biosensors and helpful discussions. We are grateful to Charlotte Sarre for the mGlu5 receptor graphic design. This work was supported by the Agence Nationale de la Recherche (JP, ANR-22-CE16-0013 LEARN) and the European Research Council (ERC) under the European Union’s Horizon 2020 research and innovation programme (JP, grant agreement No. 646788 VERTTICAL CITY).

## Author contributions

MB and JP conceived research and wrote the manuscript; MB carried out the majority of the experiments, conception, design of the protocols, analysis of the results and production of the figures. CC carried out G protein sensor experiments, including protocol design, experiments and analyses. NB performed theoretical and manual supervision of the molecular biology experiments (electrophoresis, plasmid digestion, transfection) and data interpretation. YC provided general assistance with the acquisition and analysis of results in microscopy, he also developed optimised programmes to facilitate and automate signal analysis. JFI helped to design and perform experiments to detect mGlu5 receptor conformation. EM supervised and implemented neuronal cultures with NB and the development of population BRET protocols. LT cloned the pCAGGS-TRPC6-ires2-mbtdtomato plasmid. JC took an active part in the implementation of the experimental protocol and together with LT provided advices for patch clamp recordings on HEK and analysis of results. JP carried out BRET and calcium microscopy experiments in neurons, and was involved in the analysis of results and theoretical supervision at all levels. JP conceived and supervised the full project. All authors contribute to the preparation of the manuscript and approved it.

## References

1. Lagerström MC, Schiöth HB. Erratum: Structural diversity of G protein-coupled receptors and significance for drug discovery. Nat Rev Drug Discov. 2008;7(6).

2. Ben-Chaim Y, Tour O, Dascal N, Parnas I, Parnas H. The M2 muscarinic G-protein-coupled receptor is voltage-sensitive. Journal of Biological Chemistry. 2003;278(25).

3. Ben-Chaim Y, Chanda B, Dascal N, Bezanilla F, Parnas I, Parnas H. Movement of ‘gating charge’ is coupled to ligand binding in a G-protein-coupled receptor. Nature. 2006;444(7115).

4. Rinne A, Mobarec JC, Mahaut-Smith M, Kolb P, Bünemann M. The mode of agonist binding to a G protein-coupled receptor switches the effect that voltage changes have on signaling. Sci Signal. 2015;8(401).

5. Navarro-Polanco RA, Galindo EGM, Ferrer-Villada T, Arias M, Rigby JR, Sánchez-Chapula JA, et al. Conformational changes in the M2 muscarinic receptor induced by membrane voltage and agonist binding. Journal of Physiology. 2011;589(7).

6. Martinez-Pinna J, Gurung IS, Vial C, Leon C, Gachet C, Evans RJ, et al. Direct voltage control of signaling via P2Y1 and other Gαq-coupled receptors. Journal of Biological Chemistry. 2005;280(2).

7. Ruland JG, Kirchhofer SB, Klindert S, Bailey CP, Bünemann M. Voltage modulates the effect of μ-receptor activation in a ligand-dependent manner. Br J Pharmacol. 2020;177(15).

8. Sahlholm K, Nilsson J, Marcellino D, Fuxe K, Århem P. Voltage-dependence of the human dopamine D2 receptor. Synapse. 2008;62(6).

9. Kurz M, Krett AL, Bünemann M. Voltage dependence of prostanoid receptors. Mol Pharmacol. 2020;97(4).

10. Barchad-Avitzur O, Priest MF, Dekel N, Bezanilla F, Parnas H, Ben-Chaim Y. A Novel Voltage Sensor in the Orthosteric Binding Site of the M2 Muscarinic Receptor. Biophys J. 2016;111(7).

11. Ohana L, Barchad O, Parnas I, Parnas H. The metabotropic glutamate G-protein-coupled receptors mGluR3 and mGluR1a are voltage-sensitive. Journal of Biological Chemistry. 2006;281(34).

12. Hoppe A, Marti-Solano M, Drabek M, Bünemann M, Kolb P, Rinne A. The allosteric site regulates the voltage sensitivity of muscarinic receptors. Cell Signal. 2018;42.

13. López-Serrano AL, De Jesús-Pérez JJ, Zamora-Cárdenas R, Ferrer T, Rodríguez-Menchaca AA, Tristani-Firouzi M, et al. Voltage-induced structural modifications on M2 muscarinic receptor and their functional implications when interacting with the superagonist iperoxo. Biochem Pharmacol. 2020;177.

14. Kupchik YM, Barchad-Avitzur O, Wess J, Ben-Chaim Y, Parnas I, Parnas H. A novel fast mechanism for GPCR-mediated signal transduction - Control of neurotransmitter release. Journal of Cell Biology. 2011;192(1).

15. Zhang Q, Liu B, Li Y, Yin L, Younus M, Jiang X, et al. Regulating quantal size of neurotransmitter release through a GPCR voltage sensor. Proc Natl Acad Sci U S A. 2020;117(43).

16. Rozenfeld E, Tauber M, Ben-Chaim Y, Parnas M. GPCR voltage dependence controls neuronal plasticity and behavior. Nat Commun. 2021;12(1).

17. Scheefhals N, MacGillavry HD. Functional organization of postsynaptic glutamate receptors. Vol. 91, Molecular and Cellular Neuroscience. 2018.

18. Niswender CM, Conn PJ. Metabotropic glutamate receptors: Physiology, pharmacology, and disease. Vol. 50, Annual Review of Pharmacology and Toxicology. 2010.

19. Rae MG, Martin DJ, Collingridge GL, Irving AJ. Role of Ca2+ stores in metabotropic L-glutamate receptor-mediated supralinear Ca2+ signaling in rat hippocampal neurons. Journal of Neuroscience. 2000;20(23).

20. Perroy J, Richard S, Nargeot J, Bockaert J, Fagni L. Permissive effect of voltage on mGlu 7 receptor subtype signaling in neurons. Journal of Biological Chemistry. 2002;277(2).

21. Reiner A, Levitz J. Glutamatergic Signaling in the Central Nervous System: Ionotropic and Metabotropic Receptors in Concert. Vol. 98, Neuron. 2018.

22. Seeburg PH, Burnashev N, Köhr G, Kuner T, Sprengel R, Monyer H. The NMDA Receptor Channel: Molecular Design of a Coincidence Detector. In: Proceedings of the 1993 Laurentian Hormone Conference. 1995.

23. Kwag J, Paulsen O. Gating of NMDA receptor-mediated hippocampal spike timing-dependent potentiation by mGluR5. Neuropharmacology. 2012;63(4).

24. Aniksztejn L, Otani S, Ben-Ari Y. Quisqualate Metabotropic Receptors Modulate NMDA Currents and Facilitate Induction of Long-Term Potentiation Through Protein Kinase C. European Journal of Neuroscience. 1992;4(6).

25. O’neill N, McLaughlin C, Komiyama N, Sylantyev S. Biphasic modulation of NMDA receptor function by metabotropic glutamate receptors. Journal of Neuroscience. 2018;38(46).

26. Mannaioni G, Marino MJ, Valenti O, Traynelis SF, Conn PJ. Metabotropic glutamate receptors 1 and 5 differentially regulate CA1 pyramidal cell function. Journal of Neuroscience. 2001;21(16).

27. Hu JH, Park JM, Park S, Xiao B, Dehoff MH, Kim S, et al. Homeostatic Scaling Requires Group I mGluR Activation Mediated by Homer1a. Neuron. 2010;68(6).

28. Holz A, Mülsch F, Schwarz MK, Hollmann M, Döbrössy MD, Coenen VA, et al. Enhanced mGlu5 Signaling in Excitatory Neurons Promotes Rapid Antidepressant Effects via AMPA Receptor Activation. Neuron. 2019;104(2).

29. Lepannetier S, Gualdani R, Tempesta S, Schakman O, Seghers F, Kreis A, et al. Activation of TRPC1 Channel by Metabotropic Glutamate Receptor mGluR5 Modulates Synaptic Plasticity and Spatial Working Memory. Front Cell Neurosci. 2018;12.

30. Nakahara K, Okada M, Nakanishi S. The metabotropic glutamate receptor mGluR5 induces calcium oscillations in cultured astrocytes via protein kinase C phosphorylation. J Neurochem. 1997;69(4).

31. O’Malley KL, Jong YJI, Gonchar Y, Burkhalter A, Romano C. Activation of metabotropic glutamate receptor mGlu5 on nuclear membranes mediates intranuclear Ca2+ changes in heterologous cell types and neurons. Journal of Biological Chemistry. 2003;278(30).

32. Nash MS, Schell MJ, Atkinson PJ, Johnston NR, Nahorski SR, John Challiss RA. Determinants of metabotropic glutamate receptor-5-mediated Ca 2+ and inositol 1,4,5-trisphosphate oscillation frequency: Receptor density versus agonist concentration. Journal of Biological Chemistry. 2002;277(39).

33. Gómez-Santacana X, Pittolo S, Rovira X, Lopez M, Zussy C, Dalton JAR, et al. Illuminating Phenylazopyridines to Photoswitch Metabotropic Glutamate Receptors: From the Flask to the Animals. ACS Cent Sci. 2017;3(1).

34. Moutin E, Sakkaki S, Compan V, Bouquier N, Giona F, Areias J, et al. Restoring glutamate receptosome dynamics at synapses rescues autism-like deficits in Shank3-deficient mice. Mol Psychiatry. 2021;

35. Bradley SJ, Watson JM, Challiss RAJ. Effects of positive allosteric modulators on single-cell oscillatory Ca2+ signaling initiated by the type 5 metabotropic glutamate receptor. Mol Pharmacol. 2009;76(6).

36. Bradley SJ, Langmead CJ, Watson JM, Challiss RAJ. Quantitative analysis reveals multiple mechanisms of allosteric modulation of the mGlu5 receptor in rat astroglia. Mol Pharmacol. 2011;79(5).

37. Namkung Y, Le Gouill C, Lukashova V, Kobayashi H, Hogue M, Khoury E, et al. Monitoring G protein-coupled receptor and β-arrestin trafficking in live cells using enhanced bystander BRET. Nat Commun. 2016;7.

38. Doumazane E, Scholler P, Fabre L, Zwier JM, Trinquet E, Pin JP, et al. Illuminating the activation mechanisms and allosteric properties of metabotropic glutamate receptors. Proc Natl Acad Sci U S A [Internet]. 2013/03/15. 2013;110(15):E1416–25. Available from: http://www.ncbi.nlm.nih.gov/pubmed/23487753

39. Scholler P, Moreno-Delgado D, Lecat-Guillet N, Doumazane E, Monnier C, Charrier-Savournin F, et al. HTS-compatible FRET-based conformational sensors clarify membrane receptor activation. Nat Chem Biol. 2017;13(4).

40. Avet C, Mancini A, Breton B, Gouill C Le, Hauser AS, Normand C, et al. Effector membrane translocation biosensors reveal G protein and Parrestin coupling profiles of 100 therapeutically relevant GPCRs. Elife. 2022;11.

41. Goyet E, Bouquier N, Ollendorff V, Perroy J. Fast and high resolution single-cell BRET imaging. Sci Rep [Internet]. 2016/06/16. 2016;6:28231. Available from: http://www.ncbi.nlm.nih.gov/pubmed/27302735

42. Hof T, Chaigne S, Récalde A, Sallé L, Brette F, Guinamard R. Transient receptor potential channels in cardiac health and disease. Vol. 16, Nature Reviews Cardiology. 2019.

43. Wang Q, Wang D, Shibata S, Ji T, Zhang L, Zhang R, et al. Group I metabotropic glutamate receptor activation induces TRPC6-dependent calcium influx and RhoA activation in cultured human kidney podocytes. Biochem Biophys Res Commun. 2019;511(2).

44. Nagy GA, Botond G, Borhegyi Z, Plummer NW, Freund TF, Hájos N. DAG-sensitive and Ca2+ permeable TRPC6 channels are expressed in dentate granule cells and interneurons in the hippocampal formation. Hippocampus. 2013;23(3).

45. Itsuki K, Imai Y, Hase H, Okamura Y, Inoue R, Mori MX. PLC-mediated PI(4,5)P2 hydrolysis regulates activation and inactivation of TRPC6/7 channels. Journal of General Physiology. 2014;143(2).

46. Dryer SE, Kim EY. Permeation and rectification in canonical transient receptor potential-6 (TRPC6) channels. Vol. 9, Frontiers in Physiology. 2018.

47. Lutzu S, Castillo PE. Modulation of NMDA Receptors by G-protein-coupled receptors: Role in Synaptic Transmission, Plasticity and Beyond. Vol. 456, Neuroscience. 2021.

48. Pisani A, Calabresi P, Centonze D, Bernardi G. Enhancement of NMDA responses by group I metabotropic glutamate receptor activation in striatal neurones. Br J Pharmacol. 1997;120(6).

49. Heidinger V, Manzerra P, Wang XQ, Strasser U, Yu SP, Choi DW, et al. Metabotropic glutamate receptor 1-induced upregulation of NMDA receptor current: Mediation through the Pyk2/Src-family kinase pathway in cortical neurons. Journal of Neuroscience. 2002;22(13).

50. Benquet P, Gee CE, Gerber U. Two distinct signaling pathways upregulate NMDA receptor responses via two distinct metabotropic glutamate receptor subtypes. Journal of Neuroscience. 2002;22(22).

51. Ding F, O’donnell J, Xu Q, Kang N, Goldman N, Nedergaard M. Changes in the composition of brain interstitial ions control the sleep-wake cycle. Science (1979). 2016;352(6285).

52. Chiu DN, Carter BC. Synaptic NMDA receptor activity at resting membrane potentials. Front Cell Neurosci. 2022;16.

53. Sobczyk A, Scheuss V, Svoboda K. NMDA receptor subunit-dependent [Ca2+] signaling in individual hippocampal dendritic spines. Journal of Neuroscience. 2005;25(26).

54. Nasrallah C, Cannone G, Briot J, Rottier K, Berizzi AE, Huang CY, et al. Agonists and allosteric modulators promote signaling from different metabotropic glutamate receptor 5 conformations. Cell Rep. 2021;36(9).

55. Inuzuka T, Fujioka Y, Tsuda M, Fujioka M, Satoh AO, Horiuchi K, et al. Attenuation of ligand-induced activation of angiotensin II type 1 receptor signaling by the type 2 receptor via protein kinase C. Sci Rep. 2016;6.

56. García-Sáinz JA, Martínez-Alfaro M, Romero-Avila MT, González-Espinosa C. Characterization of the AT1 angiotensin II receptor expressed in guinea pig liver. Journal of Endocrinology. 1997;154(1).

57. Rössig L, Zólyomi A, Catt KJ, Balla T. Regulation of angiotensin II-stimulated Ca2+ oscillations by Ca2+ influx mechanisms in adrenal glomerulosa cells. Journal of Biological Chemistry. 1996;271(36).

58. Bockaert J, Perroy J, Ango F. The complex formed by group i metabotropic glutamate receptor (MGLUR) and homer1a plays a central role in metaplasticity and homeostatic synaptic scaling. Vol. 41, Journal of Neuroscience. 2021.

59. Bodzęta A, Scheefhals N, MacGillavry HD. Membrane trafficking and positioning of mGluRs at presynaptic and postsynaptic sites of excitatory synapses. Vol. 200, Neuropharmacology. 2021.

60. Rasmussen R, O’Donnell J, Ding F, Nedergaard M. Interstitial ions: A key regulator of state-dependent neural activity? Vol. 193, Progress in Neurobiology. 2020.

61. Francesconi W, Cammalleri M, Sanna PP. The metabotropic glutamate receptor 5 is necessary for late-phase long-term potentiation in the hippocampal CA1 region. Brain Res. 2004;1022(1–2).

62. Neyman S, Manahan-Vaughan D. Metabotropic glutamate receptor 1 (mGluR1) and 5 (mGluR5) regulate late phases of LTP and LTD in the hippocampal CA1 region in vitro. European Journal of Neuroscience. 2008;27(6).

63. Diering GH, Nirujogi RS, Roth RH, Worley PF, Pandey A, Huganir RL. Homer1a drives homeostatic scaling-down of excitatory synapses during sleep. Science (1979). 2017;355(6324).

64. Ango F, Prézeau L, Muller T, Tu JC, Xiao B, Worley PF, et al. Agonist-independent activation of metabotropic glutamate receptors by the intracellular protein Homer. Nature. 2001;411(6840).

65. Moutin E, Raynaud F, Roger J, Pellegrino E, Homburger V, Bertaso F, et al. Dynamic remodeling of scaffold interactions in dendritic spines controls synaptic excitability. Journal of Cell Biology. 2012;198(2):251–63.

66. Guo W, Ceolin L, Collins KA, Perroy J, Huber KM. Elevated CaMKIIalpha and Hyperphosphorylation of Homer Mediate Circuit Dysfunction in a Fragile X Syndrome Mouse Model. Cell Rep [Internet]. 2015/12/17. 2015;13(10):2297–311. Available from: http://www.ncbi.nlm.nih.gov/pubmed/26670047

67. Birk A, Rinne A, Bünemann M. Membrane potential controls the efficacy of catecholamineinduced β1-Adrenoceptor activity. Journal of Biological Chemistry. 2015 Nov 6;290(45):27311–20.

68. Navarro-Polanco RA, Aréchiga-Figueroa IA, Salazar-Fajardo PD, Benavides-Haro DE, Rodríguez-Elías JC, Sachse FB, et al. Voltage sensitivity of M2 muscarinic receptors underlies the delayed rectifier-like activation of ACh-gated K+ current by choline in feline atrial myocytes. Journal of Physiology. 2013;591(17).

69. Jayant K, Hirtz JJ, Plante IJ La, Tsai DM, De Boer WDAM, Semonche A, et al. Targeted intracellular voltage recordings from dendritic spines using quantum-dot-coated nanopipettes. Nat Nanotechnol. 2017;12(4).

70. Cornejo VH, Ofer N, Yuste R. Voltage compartmentalization in dendritic spines in vivo. Science (1979). 2022;375(6576).

71. Acker CD, Hoyos E, Loew LM. EPSPs measured in proximal dendritic spines of cortical pyramidal neurons. eNeuro. 2016;3(2).

72. David D, Bentulila Z, Tauber M, Ben-Chaim Y. G Protein-Coupled Receptors Regulated by Membrane Potential. Vol. 23, International Journal of Molecular Sciences. 2022.

73. Mølck C, Harpsøe K, Gloriam DE, Clausen RP, Madsen U, Pedersen L, et al. Pharmacological characterization and modeling of the binding sites of novel 1,3-Bis(pyridinylethynyl)benzenes as metabotropic glutamate receptor 5-selective negative allosteric modulators. Mol Pharmacol. 2012;82(5).

74. Rovira X, Malhaire F, Scholler P, Rodrigo J, Gonzalez-Bulnes P, Llebaria A, et al. Overlapping binding sites drive allosteric agonism and positive cooperativity in type 4 metabotropic glutamate receptors. FASEB Journal. 2015;29(1).

75. Perroy J, Raynaud F, Homburger V, Rousset MC, Telley L, Bockaert J, et al. Direct interaction enables cross-talk between ionotropic and group I metabotropic glutamate receptors. Journal of Biological Chemistry. 2008;

76. Niwa H, Yamamura K, Miyazaki J. Efficient selection for high-expression transfectants with a novel eukaryotic vector. Gene. 1991;108(2).

77. Ricart-Ortega M, Berizzi AE, Catena J, Malhaire F, Muñoz L, Serra C, et al. Development and validation of a mass spectrometry binding assay for mGlu5 receptor. Anal Bioanal Chem. 2020;412(22).

78. Moutin E, Hemonnot AL, Seube V, Linck N, Rassendren F, Perroy J, et al. Procedures for Culturing and Genetically Manipulating Murine Hippocampal Postnatal Neurons. Front Synaptic Neurosci. 2020;12.

79. Chastagnier Y, Moutin E, Hemonnot AL, Perroy J. Image processing for bioluminescence resonance energy transfer measurement-BRET-Analyzer. Front Comput Neurosci. 2018;11.

80. Lin MZ, Schnitzer MJ. Genetically encoded indicators of neuronal activity. Vol. 19, Nature Neuroscience. 2016.

