## supplemental data for "Voltage tunes mGlu5 receptor function, impacting synaptic transmission"

### Supplementary Data

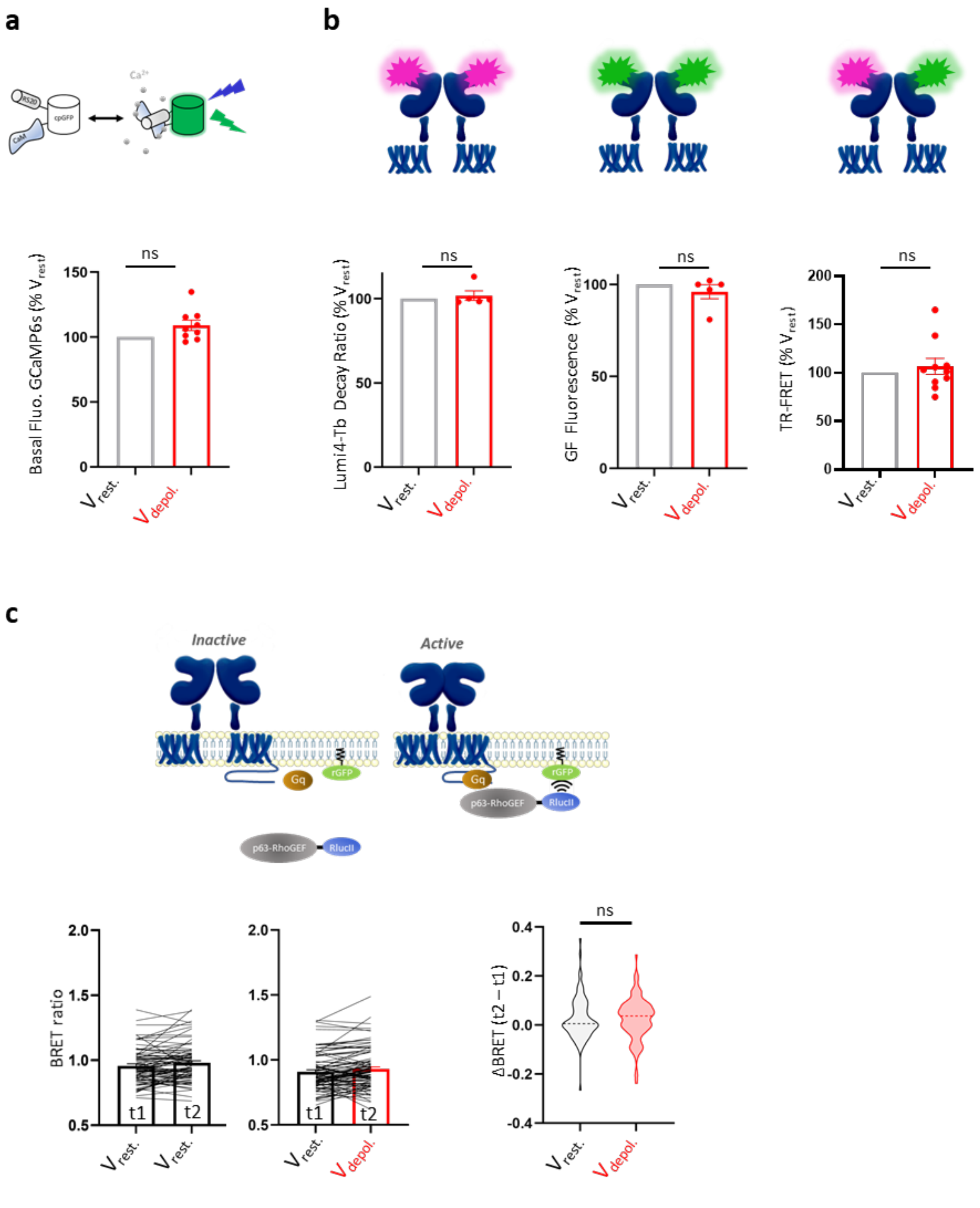


**Figure supp. 1. Lack of effect of voltage on biosensors. a)** Basal fluorescence in HEK293T cells expressing GCaMP6s and mGlu5 receptor at $V_{rest}$ and $V_{depol}$. Data are mean ± SEM of n = 9 experiments; Statistics: One sample Wilcoxon test. **b)** SNAP-Lumi4-Tb decay ratio, SNAP-Green fluorescence and TR-FRET at $V_{rest}$ and $V_{depol}$. Data are mean ± SEM of n = 4, 5 and 10 experiments; Statistics: One sample Wilcoxon test. **c)** Single-cell BRET measurements before (t1) and after (t2) medium changes from $V_{rest}$ to $V_{rest}$ (left, n = 87 cells) or $V_{depol}$ (middle, n = 97 cells). Single cell BRET t2 – t1 net signals; Statistics: Unpaired t-test.

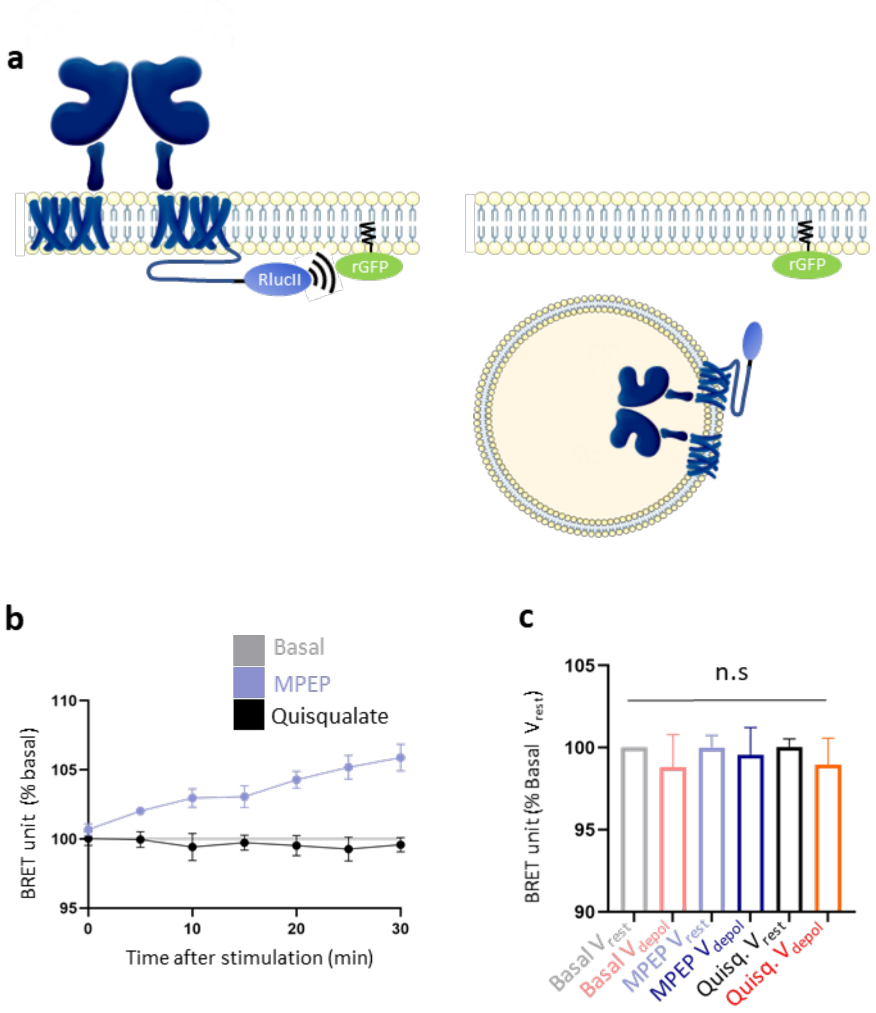


**Figure Supp. 2. Lack of effect of voltage on mGlu5 receptor membrane expression. a)** Schematic view of the ebBRET biosensor monitoring mGlu5 receptor membrane expression. **b)** BRET intensity after MPEP or quisqualate stimulation at $V_{rest}$. Each time point was normalized to the basal condition. Data are mean ± SEM of n = 3 and 4 independent experiments. **c)** BRET intensity at $V_{rest}$ and $V_{depol}$ (basal) and 2min after MPEP (10µM) or quisqualate (10µM) stimulation. Data were normalized to the basal $V_{rest}$ condition and are mean ± SEM of n = 4 independent experiments; Statistics: Kruskal-Wallis.

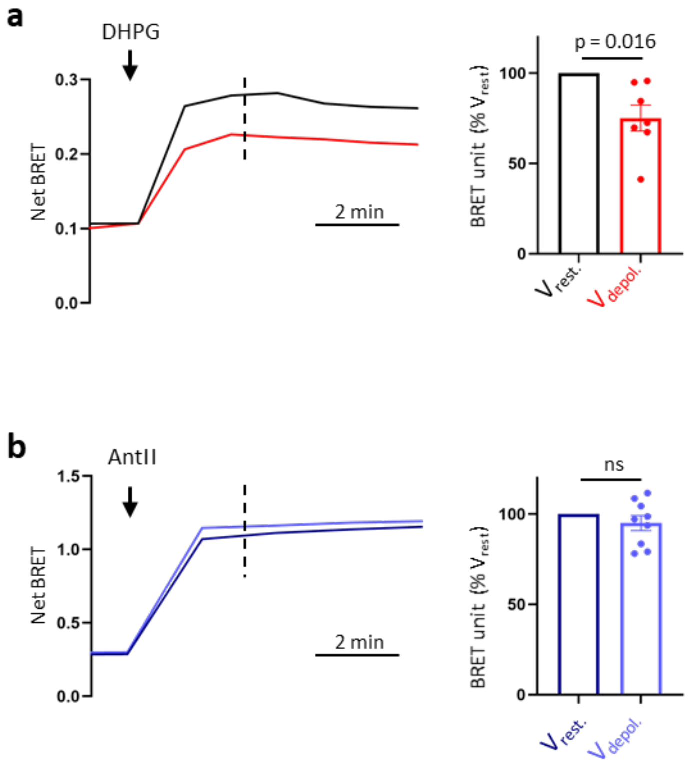


**Figure Supp. 3. Voltage tunes $G_{q/11}$ protein activation by mGlu5 receptor, but not AT1 receptor. a)** Representative kinetics of the net BRET signal after DHPG (100µM) stimulation at $V_{rest}$ or $V_{depol}$ (left) and mean ± SEM BRET signal, 2min after stimulation n = 7 independent experiments (right); Statistics: One sample Wilcoxon test. **b)** Same legend as (a), after Angiotensin II (1µM) stimulation of AT1 receptor at $V_{rest}$ or $V_{depol}$. (n = 9 independent experiments).

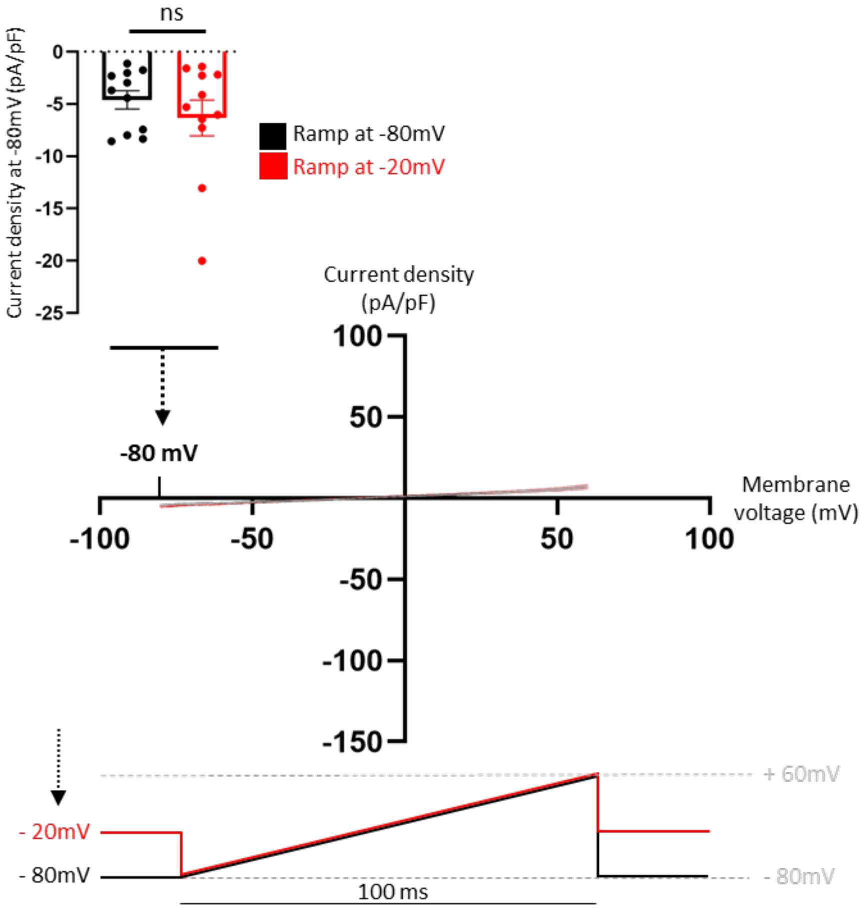


**Figure Supp. 4. Voltage does not change cell conductance.** mGlu5-Venus/TRPC6-tomato co-transfected cells were held at -80mV (black) or -20mV (red), without stimulating receptors. Current-voltage relationship was recorded for potentials ranging from -80mV to +60mV in 100ms (bottom). Inset: Mean ± SEM current density at -80mV. Data are mean ± SEM of n = 11 cells per condition from 4 independent experiments; Statistics: Mann-Whitney test.

**Video legends**

**Video S1 and S2:** live cell imaging of GCaMP6s fluorescence at $V_{rest}$ (S1) or $V_{depol}$ (S2) before and during quisqualate (10µM) application, accelerated 50 times.

**Video S3:** live cell imaging of GCaMP6s fluorescence in basal condition (ACSF containing 0.7 mM $Mg^{2+}$), followed by successive addition of DHPG (50 µM) and then AP5 (50 µM). The 3 recording sequences last for 2 min and 15 sec each, the movie is accelerated 10 times.
